# Broadly neutralizing and protective nanobodies against diverse sarbecoviruses

**DOI:** 10.1101/2022.04.12.488087

**Authors:** Mingxi Li, Yifei Ren, Zhen Qin Aw, Bo Chen, Ziqing Yang, Yuqing Lei, Lin Cheng, Qingtai Liang, Junxian Hong, Yiling Yang, Jing Chen, Yi Hao Wong, Sisi Shan, Senyan Zhang, Jiwan Ge, Ruoke Wang, Xuanling Shi, Qi Zhang, Zheng Zhang, Justin Jang Hann Chu, Xinquan Wang, Linqi Zhang

**Author notes:** These authors contributed equally (Mingxi Li, Yifei Ren, Zhen Qin Aw, Bo Chen).

## Abstract

As SARS-CoV-2 Omicron and other variants of concern continue spreading around the world, development of antibodies and vaccines to confer broad and protective activity is a global priority. Here, we report on the identification of a special group of nanobodies from immunized alpaca with exceptional breadth and potency against diverse sarbecoviruses including SARS-CoV-1, Omicron BA.1, and BA.2. Crystal structure analysis of one representative nanobody, 3-2A2-4, revealed a highly conserved epitope between the cryptic and the outer face of the receptor binding domain (RBD). The epitope is readily accessible regardless of RBD in “up” or “down” conformation and distinctive from the receptor ACE2 binding site. Passive delivery of 3-2A2-4 protected K18-hACE2 mice from infection of authentic SARS-CoV-2 Delta and Omicron. This group of nanobodies and the epitope identified should provide invaluable reference for the development of next generation antibody therapies and vaccines against wide varieties of SARS-CoV-2 infection and beyond.

## INTRODUCTION

As the severe acute respiratory syndrome coronavirus 2 (SARS-CoV-2) continues to rage around the world, we have witnessed the rapid emergence and turnover of multiple variants of concerns (VOCs) such as Alpha (B.1.1.7) initially identified in the United Kingdom; Beta (B.1.351) in South Africa; Gamma (P.1) in Brazil; Delta (B.1.617.2) in India; and Omicron (BA.1 and BA.2) in Botswana and South Africa (https://www.who.int/en/activities/tracking-SARS-CoV-2-variants/). These VOCs are not only associated with steeply increased new infections among unvaccinated but also break-through infections among the vaccinated individuals ^1–4^. Increasing evidence suggests that substantial changes in their antigenic properties have facilitated these VOCs to escape from serum neutralization of convalescent and vaccinated individuals ^5–9^. As a result, efficacies of all vaccine modalities as well as many therapeutic antibodies approved for emergency use authorization (EUA) have been severely compromised, particularly toward Omicron, followed by Beta, Delta, Gamma, and Alpha ^6, 8, 10–14^. Among these VOCs, Omicron is perhaps the most insidious as it generally causes milder symptoms among healthy and vaccinated individuals but displays exceptionally high viral load in the upper respiratory tract and extremely efficient in transmission ^15–17^. As quiet as it seems, Omicron has been actively replacing Delta and other local variants to become the most dominant variant in many parts of the world ^18^. Development of broader and more effective therapies and vaccines against Omicron and future variants has therefore become an urgent and global priority.

One striking aspect of Omicron is the largest number of mutations found in the spike (S) protein among the VOCs identified thus far (https://www.gisaid.org). It is currently unknow how Omicron accumulated such high number of mutations in such a short period, although some speculated that it could be derived from immune-compromised individuals or straight across from other animal species. BA.1 and BA.2 are the two major subvariants of Omicron that are rapidly spreading and accounting for majority of the new and break-through infections worldwide. The S protein of SARS-CoV-2 is the major target for neutralizing antibodies and has been engineered and used in many of the vaccine modalities ^19–21^. In the Omicron S protein, there are approximately 35 substitutions compared to the prototype strain found in Wuhan, China. Of which, at least 15 are located in the receptor-binding domain (RBD) and at least 8 in the N-terminal domain (NTD), although the exact number of substitutions vary among various subvariant within Omicron (https://www.gisaid.org). Recently, several elegant studies have pinpointed a few key substitutions in the S protein that are responsible for neutralization escape, and many of which are shared among the VOCs. For example, the N501Y substitution previously shown to enhance binding affinity to the receptor angiotensin-converting enzyme 2 (ACE2) is found in Alpha, Beta, Gamma, and Omicron ^22^. Delta and other variants such as Epsilon, Lambda, and Kappa have substitutions L452R or L452Q within the RBD that also facilitate virus escape ^23, 24^. Beta, Gamma, and Omicron each have three substitution sites in common within the RBD, namely K417N/T, E484K/A, and N501Y, which resulted in marked reduction or complete loss of neutralizing activities of many therapeutic antibodies and immune serum from vaccinated individuals ^5, 6, 25, 26^. In the NTD, Alpha, Beta, Gamma, and Omicron share deletions/insertions and substitutions within or near the “NTD supersite” that largely consisted of the N1 (residues 14–26), N3 (residues 141–156), and N5 (residues 246– 260) loops ^27–30^. While these findings have clearly identified several critical substitutions that confer viral escape from antibody neutralization, they also point to the critical substitutions that must overcome in order to develop broad and protective antibody therapies and vaccines against various VOCs including the most divergent Omicron BA.1 and BA.2.

We previously reported hundreds of spike-specific monoclonal antibodies (mAbs) from SARS-CoV-2 convalescent individuals ^31^. While these mAbs are being screened for broad and protective activity against Omicron and other VOCs, we decided to expand our search through immunizing alpaca, as this animal species like those in the family of *Camelidae* has a small and robust nanobody structure, allowing deeper and broader recognition of target antigens thereby to improve the likelihood of success in finding the ideal type of antibodies mentioned above. Structurally, nanobody is unique and composed of a single heavy chain with one variable domain (VHH). The smaller size (<15 kDa) and longer VHH domain facilitate their reach to targets that are otherwise inaccessible to conventional human antibodies ^32^. Their special properties in specificity, stability, thermotolerance, low immunogenicity, and ease in production in yeast and other cost-effective systems have made the nanobodies the ideal candidate for therapeutic and prophylactic interventions against SARS-CoV-2 infection ^33–46^. Through screening nanobody library from an immunized alpaca, we have identified a special group of nanobodies with exceptional breadth and potency against diverse sarbecoviruses including Omicron BA.1 and BA.2, SARS-CoV-1, as well as the key representatives from bats and pangolins coronaviruses. The IC50 reached as low as 0.042 μg/mL or 0.550 nM against WT D614G and remained relatively stable across the entire panel of the tested viruses. Although neutralization assays differ, this suggests they are amongst the most broad and potent nanobodies described to date. Passive delivery of one of the representative nanobodies, 3-2A2-4, protected K18-hACE2 mice from infection of authentic SARS-CoV-2 Delta and Omicron. Structural analysis revealed 3-2A2-4 targeted a highly conserved epitope between the cryptic and outer face of RBD, distinctive from the ACE2 binding site. This epitope is readily accessible regardless of RBD in “up” or “down” conformations. These results clearly indicate that we have identified a broadly neutralizing and protective nanobody that recognized a highly conserved epitope among a diverse panel of sarbecoviruses. The nanobody and the epitope identified should provide invaluable reference for the development of next generation antibody therapies and vaccines against wide varieties of SARS-CoV-2 infection and beyond.

## RESULTS

### Cross-neutralizing SARS-CoV-2 and SARS-CoV-1 nanobodies

We started with selection for cross-neutralizing SARS-CoV-2 and SARS-CoV-1 nanobodies, as these nanobodies would be expected to have higher probability to cross-react with other members of sarbecoviruses. To this end, we screened a yeast display VHH library constructed from the PBMCs of a serially immunized alpaca with RBD and spike of prototype SARS-CoV-2 (see Materials and Methods). Through iterative process of FACS-sorting, enriching, and finally expressing in the recombinant form with human IgG1 Fc fragment, we identified a total of 593 nanobodies capable of binding to the recombinant spike trimer of prototype SARS-CoV-2. Of which, 124 showed neutralizing activity to prototype SARS-CoV-2 pseudovirus with IC50 ranging from 0.003 to 5.399 μg/mL or 0.039 to 71.039 nM (Figure 1). Among these, 91 demonstrated cross-neutralizing activity to SARS-CoV-1, and all but two (No. 43 and 55) strongly bound to RBD (Figure. 1). Phylogenetically, these cross-neutralizing nanobodies were segregated into four major clusters (***a***, ***b***, ***c***, and ***d***) (Figure 1, red in the left panel). Clusters ***a*** and ***d*** nanobodies had equivalent average IC50 to SARS-CoV-2 and SARS-CoV-1 (0.077 vs. 0.056 μg/mL or 1.013 vs. 0.737 nM) while clusters ***b*** and ***c,*** however, demonstrated stronger activity to SARS-CoV-1 than to SARS-CoV-2 (0.036 vs. 0.499 μg/mL or 0.474 vs 6.566 nM). This suggests that the epitopes recognized by clusters ***b*** and ***c*** nanobodies could be more exposed and/or easily accessible on the SARS-CoV-1 spike.

**Figure 1.**
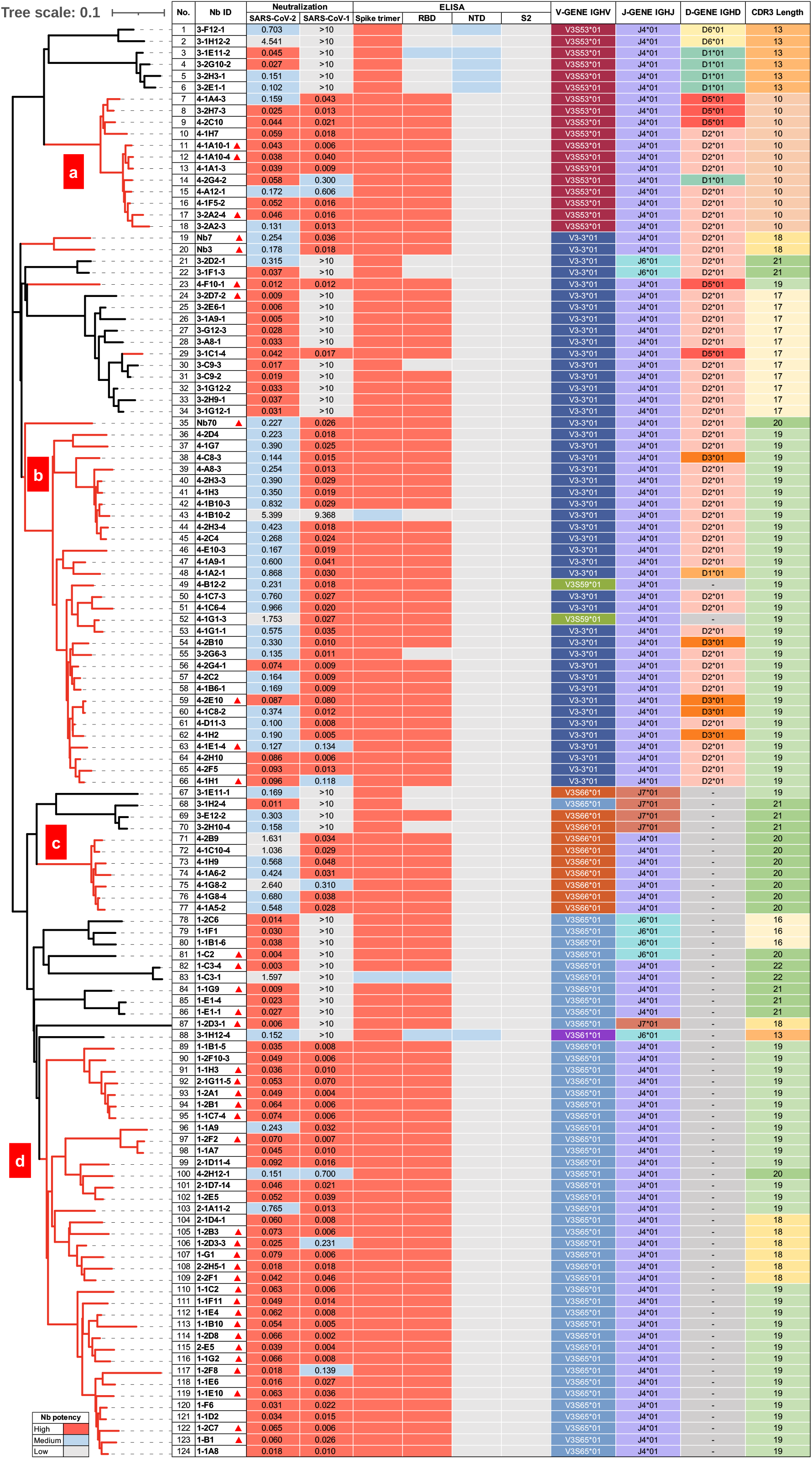
Phylogenetic, neutralizing, binding, and genetic properties of isolated nanobodies against SARS-CoV-2 and SARS-CoV-1. Phylogenetic analysis of 124 neutralizing nanobodies isolated from the VHH library of an immunized alpaca. Cross-neutralizing nanobodies against SARS-CoV-2 and SARS-CoV-1 are primarily clustered in four groups (a, b, c, and d) and highlighted by the red branches. Neutralizing activities of each nanobody to pseudoviruses carrying the spike of either prototype SARS-CoV-2 or SARS-CoV-1 are shown by half-maximal inhibitory concentration IC50 (μg/mL). The thirty-eight representative nanobodies selected for further evaluation are indicated by red triangle in Nb ID column. Results were calculated from three independent experiments and each performed in technical duplicates. The binding activity of each nanobody to prototype spike trimer, RBD, NTD, and S2 regions are indicated by the color scheme, with red for high binding, blue for medium binding, and grey for low binding. The genetic features such as germline variable gene segment (V), diversity gene segment (D), and junction gene (J) as well as CDR3 length of each nanobody are indicated with various colors. See also Figure S1.

Genetic analysis of the 91 cross-neutralizing nanobodies identified preferred usage of several germline V-gene and J-gene, particularly for IGHV3S65 (39.6%), IGHV3-3 (37.4%), IGHV3S53 (13.2%), and IGHJ4 (100%) (Figure S1a). Similar pattern of preference was also noticed among the total of 124 isolated nanobodies (Figure S1b), and the 237 published nanobodies in the CoV-AbDab database except for IGHV3S65 (Figure S1c). Such convergence on germline gene usage may suggest that the combinations of the gene segments encode nanobodies with unique structural and biochemical properties rendering them naturally complementary in shape and strong in binding to the spike surface of SARS-CoV-2 and SARS-CoV-1. Furthermore, the 91 cross-neutralizing nanobodies as well as the isolated 124 nanobodies were all dominated by 19-residue long CDR3 (Figure S1d and S1e) while those in the CoV-AbDab by 17- or longer segments (Figure S1f). However, the CDR3 length per se was unlikely a prerequisite for neutralization breadth across SARS-CoV-2 and SARS-CoV-1. Nanobodies in clusters a and d had an average CDR3 length of 10 and 19 residues, respectively, but demonstrated equivalent cross-neutralizing activity against SARS-CoV-2 and SARS-CoV-1 (Figure 1). Interestingly, sequence logo plots identified 7 polar residues (GSYYYCS) in the 19-residue CDR3 that were rather prevalent among the 91 cross-neutralizing, the 124 isolated nanobodies and the 237 published nanobodies in the CoV-AbDab database, although no obvious patterns were found in 17- or 18-residue CDR3 sequences (Figure S1g, S1h, and S1i).

### Broadly neutralizing sarbecoviruses nanobodies

To identify nanobodies that have broader activity beyond prototype SARS-CoV-2 and SARS-CoV-1, we selected 32 cross-neutralizing nanobodies based on their representation in neutralizing potency and locations on the phylogenetic tree. Six SARS-CoV-2-specific neutralizing nanobodies were also selected and used as controls. These selected nanobodies are indicated by the red triangles in Figure 1. We first classified the 32 cross-neutralizing nanobodies into 3 groups based on their degree of competition with ACE2 and VHH72, a previously published nanobody with known epitope on the cryptic face of RBD that only became accessible when RBD in the “up” standing conformation ^35^. As shown in Figure 2a, the Group 1 (G1) nanobodies strongly competed with both ACE2 and VHH72 for binding to RBD, suggesting they also bound to the similar cryptic epitopes on RBD from the angles that restricted ACE2 binding. The Group 2 (G2) nanobodies moderately competed with ACE2 but strongly with VHH72, indicating similar cryptic epitopes like those in G1 but less interfering with ACE2 binding. The Group 3 (G3) nanobodies, however, moderately competed with ACE2 or VHH72, and their epitopes and binding poses must deviate away from that of ACE2 and VHH72. The Group 4 (G4) control nanobodies included the six SARS-CoV-2 only neutralizing nanobodies, and showed varying competition activity with ACE2 and the weakest with VHH72 among all nanobodies studied here (Figure 2a). We next studied the neutralizing breadth of 38 nanobodies against a panel of 18 sarbecoviruses that dependent on ACE2 for entry, including 13 major SARS-CoV-2 variants, SARS-CoV-1, and 4 representatives from bats and pangolins. These nanobodies demonstrated diverse and distinctive neutralizing patterns, corresponding well to the 4 groups defined by the competition assays (Figure 2a, Figures S2 and S3). The G1 nanobodies showed broad neutralizing activity against all sarbecoviruses tested with average IC50 ranging from 0.085 μg/mL or 1.114 nM against WT D614G. The IC50s to Omicron BA.1 and BA.2 dropped below the detection limit (BDL in dark red) against Omicron BA.1 and BA.2 while remained relatively unchanged to other SARS-CoV-2 variants (in white), relative to WT D614G (Figure 2a, and Figure S2). Interestingly, many nanobodies showed improved neutralizing activity to SARS-CoV-1, pangolin coronavirus GD, and bat coronavirus RaTG13 (in blue) while varied considerably to pangolin coronavirus GX and bat coronavirus WIV16 (mixed with red and blue) (Figure 2a, and Figure S3). The G2 nanobodies demonstrated relatively weaker neutralizing activity against WT D614G than those in the G1 with average IC50 ranging from 0.204 μg/mL or 2.690 nM against WT D614G. However, they appeared to be less affected by BA.1. Nb70 could also neutralize BA.2 although with marked reduced activity. Subtle differences may exist in the epitope specificity and binding pose between the G1 and G2 nanobodies. Like those in the G1, the G2 nanobodies also demonstrated improved neutralizing activity to SARS-CoV-1, and those representative coronaviruses from bats and pangolins (Figure 2a). The G3 nanobodies exhibited the broadest neutralizing activity against all sarbecoviruses tested with average IC50 ranging from 0.040 μg/mL or 0.530 nM against WT D614G, although partial reduction was found to pangolin coronavirus GX. The G3 nanobodies therefore may recognize conserved epitopes on RBD distinctive from the ACE2 and VHH72 binding sites. The G4 nanobodies, despite being the strongest against WT D614G with average IC50 ranging from 0.014 μg/mL or 0.184 nM against WT D614G, had the poorest breadth against the 18 sarbecoviruses tested. Severe reduction or complete loss of neutralizing activities were found in many nanobodies with only exception to SARS-CoV-2 variants (Alpha, Epsilon, and A23.1) and pangolin coronavirus GD. As only minor variation existed among these viruses in the receptor-binding motif (RBM) (https://www.gisaid.org), it was likely that the G4 nanobodies targeted sites on RBD that either overlapped with or proximate to RBM, consistent with their noticeable competition with ACE2 for binding to RBD (Figure 2a).

**Figure 2.**
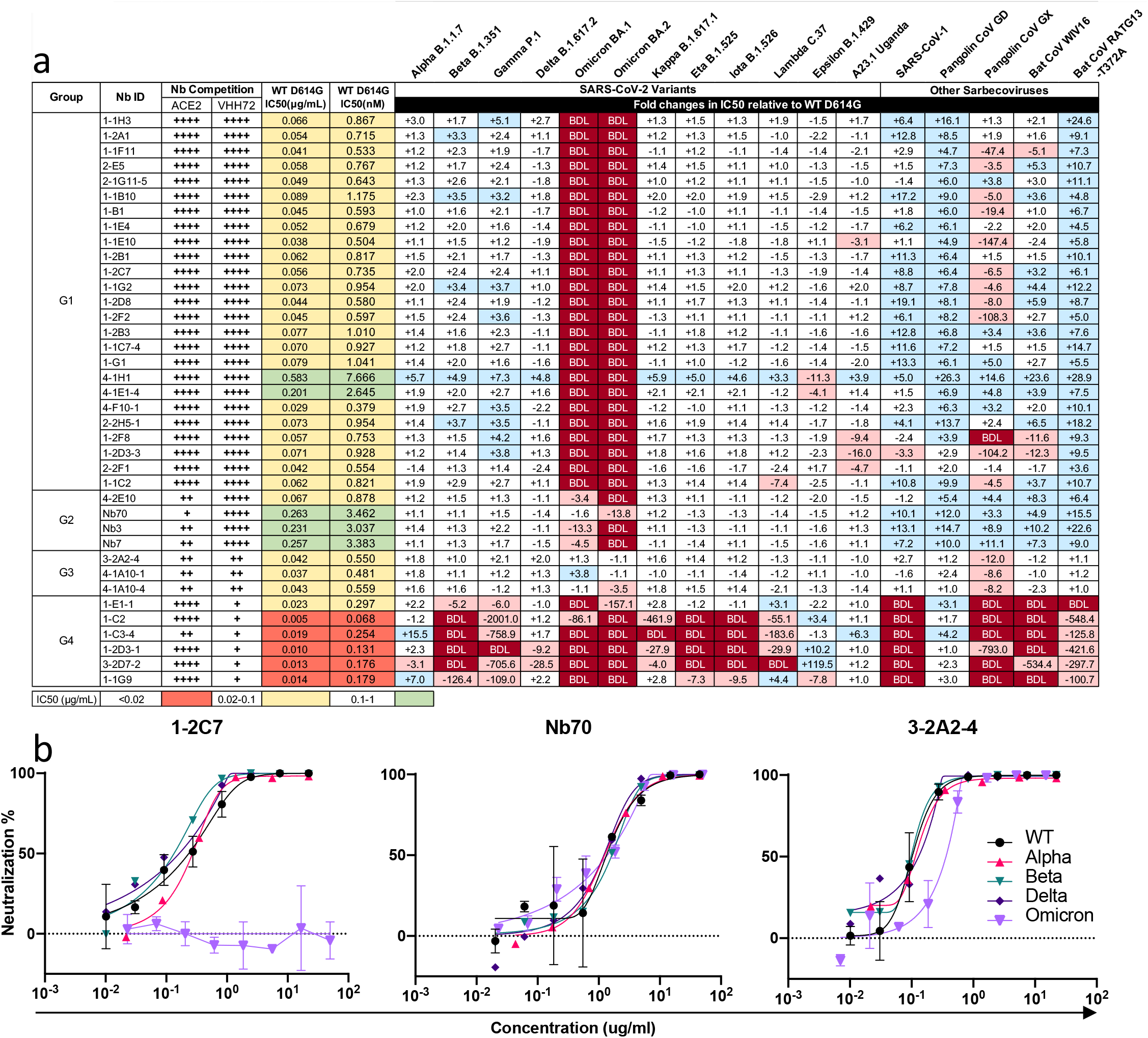
Classification and neutralizing activity of isolated nanobodies against SARS-CoV-2 variants and hACE2-dependent sarbecoviruses. **(a)** Classification of nanobodies into four major groups based on their degrees of competition with ACE2 and control VHH72 nanobody with known epitope specificity, measured by surface plasmon resonance (SPR). “++++” indicates >90% competition; “++” 30-60%, and “+” 10-30%. Neutralizing activities of nanobodies against WT D614G were presented by the actual values of 50% inhibitory concentration (IC50) while that against SARS-CoV-2 variants and hACE2-dependent sarbecoviruses were in fold-changes relative to that of WT D614G. IC50 values highlighted in salmon indicate < 0.02 μg/mL; in yellow 0.02-0.1 μg/mL, and in green >0.1 μg/mL. ‘‘-’’ indicates increased resistance and ‘‘+’’ indicates increased sensitivity to nanobody neutralization. The fold-changes highlighted in red indicate that resistance increased at least 3-fold; in blue, sensitivity increased at least threefold; and in white, resistance or sensitivity increased less than 3-fold. BDL indicates that nanobodies at their highest concentration (13.33 μg/mL) failed to reach 50% neutralization. Results were calculated from three independent experiments and each performed in technical duplicates. **(b)** Neutralizing activity of representative nanobodies from G1, G2, and G3 against authentic SARS-CoV-2 of wildtype (WT) and various VOCs such as Alpha, Beta, Delta and Omicron. Error bars indicate standard error of mean between technical replicates. See also Figure S2 and S3.

We then selected one nanobody each from G1 (1-2C7), G2 (Nb70), and G3 (3-2A2-4) to test their neutralizing activity against authentic SARS-CoV-2 such as the wildtype (WT), Alpha, Beta, Delta and Omicron variants, using focus reduction neutralization test (FRNT). Consistent with their respective activities to pseudoviruses, 3-2A2-4 was the most broad and potent among the three nanobodies with 0.102 μg/mL against WT, 0.115 μg/mL against Alpha, 0.098 μg/mL against Beta, 0.130 μg/mL against Delta, and 0.360 μg/mL against Omicron BA.1 (Figure 2b). Nb70 had a relatively moderate IC50 values with 1.337 μg/mL against WT, 1.242 μg/mL against Alpha, 1.635 μg/mL against Beta, 1.210 μg/mL against Delta, and 1.381 μg/mL against Omicron BA.1. However, 1-2C7 had IC50 values of 0.234, 0.270, 0.134, and 0.163 μg/mL against WT, Alpha, Beta, and Delta, respectively, but failed to neutralize BA.1 at the highest concentration (Figure 1b).

### Structural definition of three nanobodies epitopes

We determined crystal structures of 1-2C7, Nb70, and 3-2A2-4 bound to various recombinant RBDs (Figure 3). Of which, 1-2C7 bound to the SARS-CoV-2 Beta RBD (SARS-CoV-2 SA-RBD) was resolved at 1.8 Å resolution (Figure 3a and Table S1) whereas 3-2A2-4 bound to the SARS-CoV-2 wild type RBD (SARS-CoV-2 WT-RBD) at 2.4 Å resolution (Figure 3d and Table S1). The crystal structure of Nb70 was determined in two forms. One was the ternary complex of Nb70 and human antibody P2C-1F11 simultaneously bound to the SARS-CoV-2 WT-RBD at 2.4 Å resolution (Figure 3b and Table S1). The other was the binary complex of Nb70 bound to the SARS-CoV-1 wild type RBD (SARS CoV-1 WT-RBD) at 2.4 Å resolution (Figure 3c and Table S1).

**Figure 3.**
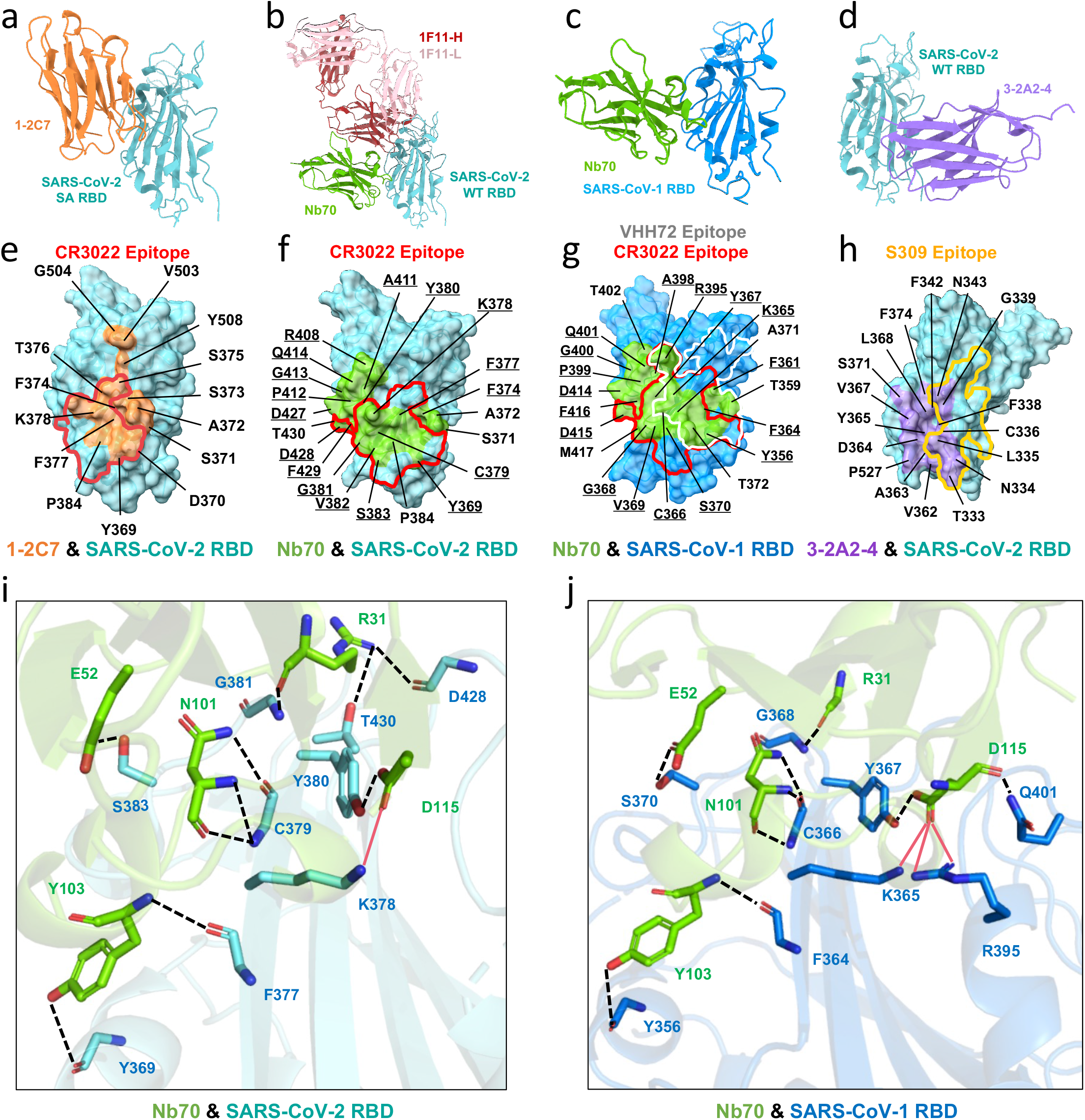
Crystal structures and epitope of three representative nanobodies bound to SARS-CoV-2 RBD or SARS-CoV-1 RBD. **(a, d)** Crystal structures of 1-2C7 and 3-2A2-4 bound to SARS-CoV-2 RBD. **(b)** Crystal structure of Nb70 and 1F11 bound to SARS-CoV-2 RBD. SARS-CoV-2 RBD is colored in cyan whereas 1-2C7 in orange, Nb70 in green, and 3-2A2-4 in purple. **(c)** Crystal structure of Nb70 bound to SARS-CoV-1 RBD. Nb70 is shown in green while the SARS-CoV-1 RBD in blue. **(e, h)** The epitope of 1-2C7 (orange) or of 3-2A2-4 (purple) is respectively depicted on the surface of SARS-CoV-2 RBD. **(f)** The epitope of Nb70 (green) depicted on the surface of SARS-CoV-2 RBD or **(g)** on the prototype SARS-CoV-1 RBD. The same residues between SARS-CoV-2 and SARS-CoV-1 epitopes were underlined. The epitope of CR3022 highlighted in red (PDB: 6YM0 on SARS-CoV-2 RBD and PDB: 7JN5 on SARS-CoV-1 RBD) is superposed onto that of **(e)** 1-2C7, **(f)** Nb70, as well as **(g)** Nb70 together with VHH72 epitope in white (PDB: 6WAQ). The epitope of S309 in yellow (PDB: 7R6W) is superimposed onto **(h)** that of 3-2A2-4. **(i, j)** Conserved interactions between Nb70 and SARS-CoV-2 RBD or SARS-CoV-1 RBD. See also Figure S4.

The crystal structures showed that both 1-2C7 and Nb70 bound to the cryptic face of SARS-CoV-2 RBD, in complete agreement with the competition data shown in Figure 2a. Their epitopes substantially overlapped with the cryptic epitopes recognized by VHH72 and the typical Class 4 antibody CR3022 (Figures 3e, 3f, and 3g). Seven residues in the 1-2C7 epitope (Y369, N370, S375, T376, F377, K378 and P384) and nine residues in the Nb70 epitope (Y369, F377, K378, C379, Y380, G381, V382, S383 and P384) were also part of the CR3022 epitope ^47^. However, 1-2C7 epitope deviated upward from the CR3022 epitope and therefore expected to create steric hindrance to ACE2 (Figure 3e). Nb70 epitope, on the other hand, was largely confined to the lower part of RBD cryptic face and further away from the ACE2 binding site (Figure 3f). In addition, the Nb70 epitopes were highly conserved between SARS-CoV-2 and SARS-CoV-1 (Figure 3f and 3g). A total of 17 residues were shared between the two respective epitopes, namely Y369, F374, F377, K378, C379, Y380, G381, V382, S383, R408, A411, P412, G413, Q414, D427, D428, and F429 on the SARS-CoV-2 RBD and Y356, F361, F364, K365, C366, Y367, G368, V369, S370, R395, A398, P399, G400, Q401, D414, D415, and F416 on the SARS-CoV-1 RBD (Figure 3f and 3g). These residues interacted with R31, E52, N101, Y103 and D115 of Nb70 through hydrogen bonds, salt-bridges, and van der Waals contacts (Figure 3i and 3j, Table S2). For instance, D115 formed hydrogen bonds and salt bridges with K378 and Y380 at the Nb70-SARS-CoV-2 interface and with K365, Y367, R395 and Q401 at the Nb70-SARS-CoV-1 interface (Figure 3i and 3j, Table S2). Such conserved epitopes provided structural basis for the cross-neutralizing activity of Nb70 between SARS-CoV-2 and SARS-CoV-1. On the other hand, 3-2A2-4 bound to the bottom part of RBD core subdomain between the cryptic and the outer face, distinctive from that of 1-2C7 and Nb70 (Figure 3h). Five residues (T333, N334, L335, G339 and N343) within the 3-2A2-4 epitope overlapped with that of the Class 3 antibody S309 (Figure 3h). Such binding pose was not expected to have steric clash with either ACE2 or VHH72, compatible with the competition results (Figure 2a).

The binding of 1-2C7, Nb70 and 3-2A2-4 to the trimeric spike is expected to be influenced by the degree of exposure and accessibility of their epitopes when RBD fluctuates between the “up” and “down” conformation. By superimposing the RBD-nanobody crystal structures onto the spike cryo-EM structures (Figure S4), we found that 1-2C7 and Nb70 could bind to the spike trimer only when two or three RBDs in the “up” state where sufficient space became available for accessing the cryptic domain on the inner surface of RBD. By contrast, 3-2A2-4 was able to bind to the spike trimer regardless of RBD in either the “up” or “down” conformation, suggesting its epitope was constantly exposed and readily accessible by the nanobody.

### 3-2A2-4 nanobody maintains neutralizing activity to Omicron BA.1 and BA.2 subvariants

We next studied which mutations found in the Omicron variant potentially responsible for the varying impact on neutralizing activity of 1-2C7, Nb70, and 3-2A2-4. Based on the structural information on the epitopes, we focused on six single (G339D, S371L, S371F, S373P, S375F, and T376A) and one triple (S371L/S373P/S375F) substitutions found in Omicron variant that were either within or in close proximity to the respective epitopes (Figure 4a-4d). Pseudoviruses bearing these single and triple mutations were constructed and tested against a serial dilution of 1-2C7, Nb70, and 3-2A2-4 (Figure 4e-4g). As single S375F substitution failed to mediate detectable infection despite multiple attempts, this particular mutant was removed from the subsequent studies. Among the viable mutants tested, 1-2C7 was mostly affected by single S371F substitution, followed by S371L, S373P, and then G339D. The triple mutations S371L/S373P/S375F resulted in complete loss of activity to the degree that was compatible to Omicron subvariants BA. 1 and BA. 2 (Figure 4e). Nb70, however, was only mildly affected by Omicron BA.1 and moderately by BA.2 (Figure 4f). Single S371 substitution led to marked reduction (S371L) or completed loss (S371F) of Nb70 activity, but the remaining single (G339D and S373P) or triple (S371L/S373P/S375F) substitutions only had moderate or no effect at all (Figure 4f). It was possible that the constellation of S371L/S373P/S375F substitutions somehow restored the epitope structure disrupted by the single S371L substitution, allowing Nb70 to resume binding and exerting neutralizing activity to Omicron BA.1. However, S371F was responsible for dramatic reduction against Omicron BA. 2 (Figure 4c). By contrast, 3-2A2-4 was the most resilient to Omicron variant among the three nanobodies tested (Figure 4g). 3-2A2-4 remained similar neutralizing activity to BA.1, BA. 2 and WT D614G, with IC50 of 0.032 μg/mL, 0.047 μg/mL, and 0.043 μg/mL respectively. Single substitutions such as G339D, S371L, or S373P had only moderate effect while the triple S371L/S373P/S375F substitutions, like that occurred to Nb70, restored neutralizing activity indistinguishable to that of WT D614G (Figure 4g). Lastly, sequence alignment of RBD sequences from the multiple sarbecoviruses revealed that epitope residues of 3-2A2-4 and Nb70 were more conserved than that of 1-2C7 (data not shown), providing molecular basis for the broad and potent neutralizing activity of 3-2A2-4 against all sarbecoviruses tested including Omicron BA.1 and BA.2.

**Figure 4.**
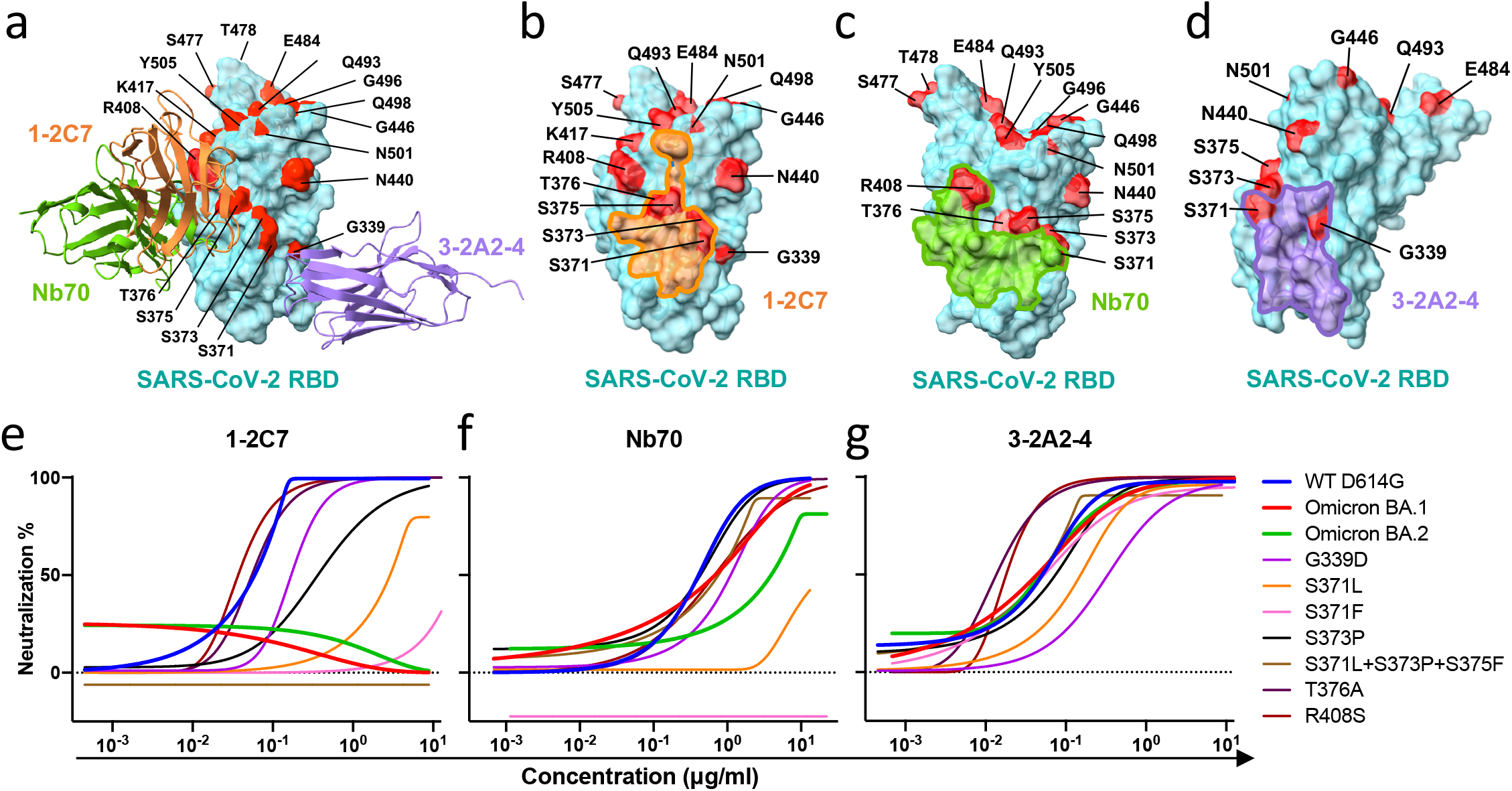
Impact of Omicron mutations on neutralizing activity of the three representative nanobodies. **(a)** Binding modes of Nb70 (green), 1-2C7 (orange) and 3-2A2-4 (purple) to prototype SARS-CoV-2 RBD with the relevant major mutations found in Omicron highlighted in red. **(b, c, d)** The footprints of 1-2C7 (orange), Nb70 (green), and 3-2A2-4 (purple) on the surface of prototype SARS-CoV-2 RBD (cyan), relative to relevant major mutations found in Omicron highlighted in red. **(e, f, g)** Neutralizing activity of 1-2C7, Nb70, and 3-2A2-4 against pseudoviruses bearing the indicated mutations found in Omicron. Data are presented as the means ± SEM from three independent experiments.

### 3-2A2-4 protects K18-hACE2 mice from infection with authentic SARS-CoV-2 Omicron and Delta

We next studied the protective potential of 3-2A2-4 against infection of authentic Omicron and Delta variants in a K18-hACE2 mouse model of SARS-CoV-2 infection, as previously described (Figure. 5a) ^48^. Specifically, the mice were intraperitoneally administered with 3-2A2-4 at a dose of 10 mg/kg body weight 24 hours prior to intranasal challenge with 1.7×10^3^ plaque-forming units (PFU) of authentic SARS-CoV-2 Omicron or 10^3^ PFU of SARS-CoV-2 Delta. The animals were then monitored daily throughout the following 14 days for their body weight and survival. Half of the animals were euthanized on day 3 post inoculation to obtain lung tissues for viral load and histopathological analysis.

**Figure 5.**
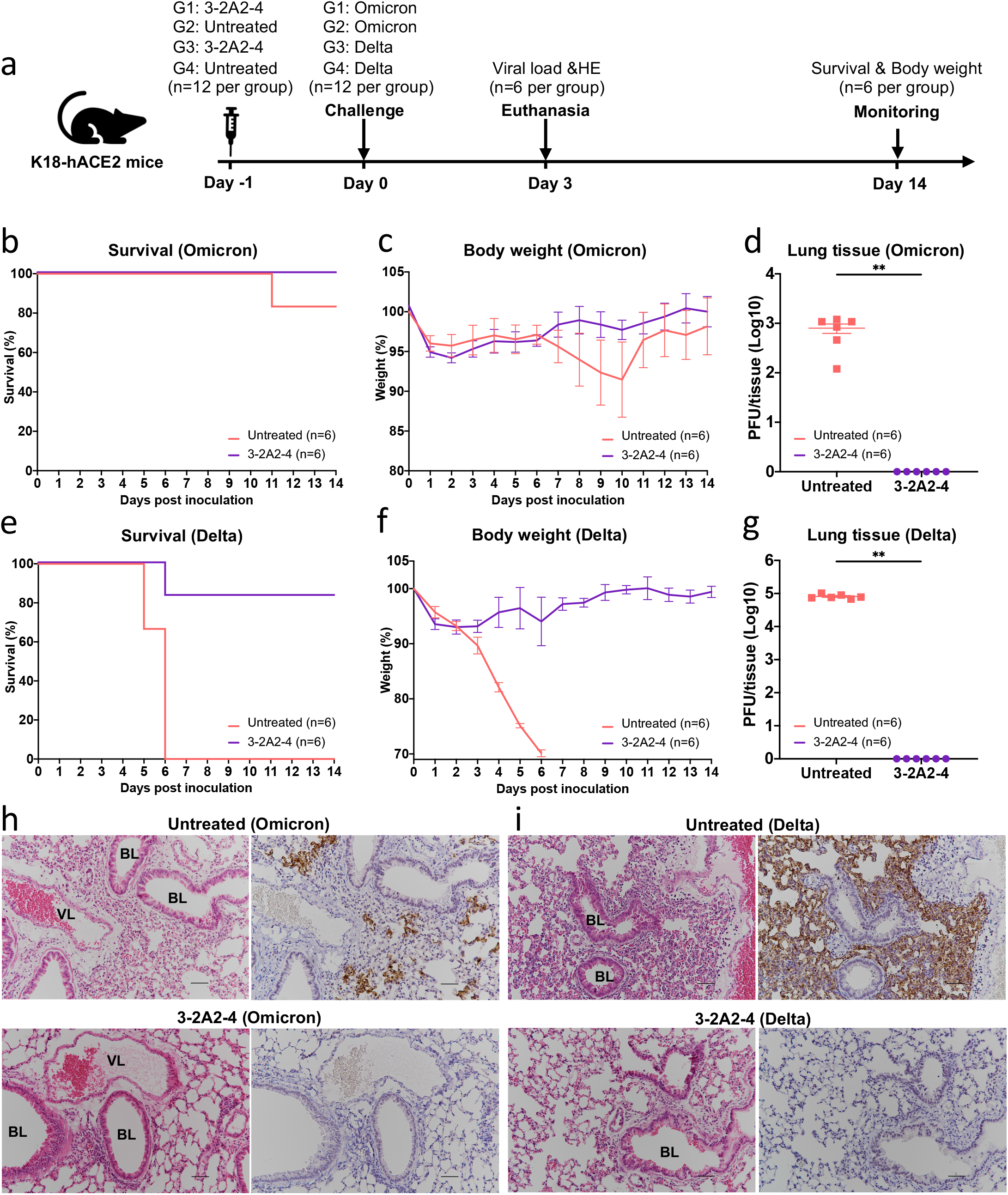
Efficacy of 3-2A2-4 prophylaxis against the authentic SARS-CoV-2 Omicron and Delta in K18-hACE-2 mice. **(a)** Experimental schedule for nanobody prophylaxis. Eight-week-old K18-hACE2 transgenic female mice were administered with 10 mg/kg body weight of 3-2A2-4 intraperitoneally or untreated 1 day prior to challenge with 1.7×10^3^ plaque-forming units (PFU) infectious SARS-CoV-2 Omicron or 10^3^ PFU Delta via the intranasal route. The survival percentage **(b** and **e)** and body weight **(c** and **f)** were recorded daily after infection until the occurrence of death or until the end of experiment. The viral load in the lung tissue **(d** and **g)** was tested by plaque assays in the tissue homogenates at 3 days post inoculation. Data are presented as the means ± SEM. Analysis of Mann-Whitney test was used. **P<0.01. **(h** and **i)** HE and IHC staining of lung tissue from 3-2A2-4-treated or untreated mice at 3 days post inoculation. VL, vascular lumen; BL, bronchiolar lumen. Scale bars, 50 μm. Each image is representative of each group.

In SARS-CoV-2 Omicron challenged groups, one out of the six untreated animals succumbed to disease on day 11 post infection whereas all 3-2A2-4 treated mice remained healthy and survived infection (Figure 5b). The changes in body weight followed the similar trend of survival, with moderate loss in untreated compared to relative stability in 3-2A2-4 treated animals, although animals in both groups experienced minor body weight loss during the first 6 days after challenge (Figure 5c). No detectable levels of live viruses were found in the lungs of 3-2A2-4-treated mice on day 3 post challenge while that in untreated mice reached an average as high as 796.7 PFU/tissue (Figure. 5d). Immunohistochemistry analysis revealed that the lung tissue of 3-2A2-4-treated mice remained intact and no viral antigen positive cells were detected (Figure. 5h). By contrast, the lung sections of untreated mice presented moderate damage and inflammation with marked infiltration of inflammatory cells. Infected cells were readily detectable using anti-N protein specific antibody (Figure. 5h).

In SARS-CoV-2 Delta challenged groups, untreated animals exhibited faster disease progression and greater severity compared to those in the Omicron challenged group, indicating Delta was more pathogenic than Omicron in this model of SARS-CoV-2 infection, in complete agreement with those recently reported ^15, 16^. All untreated animals succumbed to diseases by day 6 after challenge, and associated with severe body weight loss (Figure 5e and 5f). By contrast, 3-2A2-4-treated group remained fully protected and maintained stable body weights, except for one mouse had to be euthanized on day 6 after infection due to the requirement of experimental protocol when body weight fell below 75% of baseline (Figure 5e and 5f). No detectable levels of viruses in the lungs were found in 3-2A2-4-treated mice while that in untreated mice reached as high as 8.1×104 PFU/tissue on average (Figure 5g). The lung sections from untreated animals showed severe lung damage and inflammation with marked infiltration of inflammatory cells. A large number of viral antigen positive pneumocytes were clearly visible (Figure 5i). By contrast, in 3-2A2-4-treated mice, lung tissue remained relatively intact and well-defined. Collectively, these results indicated that the broad and potent neutralizing nanobody 3-2A2-4 conferred strong protection in vivo against challenge of authentic SARS-CoV-2 Omicron and Delta.

## DISCUSSION

The rapid emergence and spread of antigenically distinct SARS-CoV-2 variants such as Omicron BA.1 and BA.2 have resulted in the substantial reduction and loss of activity of many therapeutic antibodies and vaccines ^8,9,11–13^. Given the unusual huge number of mutations found in the spike of Omicron BA.1 and BA.2, one of the urgent questions need to be addressed is whether conserved and protective epitopes still exist on the spike trimer that can be targeted for the development of broad and potent antibody therapies and vaccines. Here, we report on the isolation and characterization of a unique group of nanobodies from immunized alpaca. The most outstanding feature of these nanobodies, exemplified by 3-2A2-4, is their exceptional breadth and potency against a highly diverse panel of sarbecoviruses including Omicron BA.1 and BA.2, SARS-CoV-1, and the key representatives from bats and pangolins coronaviruses. The IC50 reached as low as 0.042 μg/mL or 0.550 nM against WT D614G and remained relatively stable across the entire panel of the tested viruses. Although neutralization assays differ, this suggests they are amongst the broadest and most potent nanobodies described to date ^33–46^. Passive delivery of 3-2A2-4 protected K18-hACE2 mice from infection of two widely spreading SARS-CoV-2 variants Delta and Omicron. Crystal structure analysis together with modeling in the context of spike trimer revealed 3-2A2-4 recognized a highly conserved epitope between the cryptic and outer face of RBD, distinctive from the ACE2 binding site, and readily accessible in both “up” or “down” conformations. Such unique binding pose and specificity provide structural basis for 3-2A2-4 and the other members of the G3 nanobodies in withstanding the mutant residues commonly found in the major variants that compromised many therapeutic antibodies approved for EUA ^8, 12^. These results clearly indicated that we have identified a broad and protective nanobody that recognized a highly conserved epitope among a diverse panel of sarbecoviruses including Omicron BA.1 and BA.2. The nanobody and the epitope it recognized may serve as an important reference for the development of next generation antibody therapies and vaccines against wide varieties of SARS-CoV-2 infection and beyond.

Owing to their nanoscale and extended CDR3, nanobodies are expected to penetrate deeper into the spike trimer and access to hidden epitopes that are less frequently exposed and/or unreachable by conventional antibodies. This is particularly true for the cross-neutralizing SARS-CoV-2 and SARS-CoV-1 nanobodies characterized here. Virtually all nanobodies in the G1 and G2 were able to compete with a published nanobody VHH72 known to recognize the hidden epitope accessible only when RBD is in the up conformation ^35^. By contrast, the nanobodies in the G3 bound to the epitopes that are readily accessible regardless of up or down conformations of RBD. Interestingly, by screening and characterizing hundreds of monoclonal antibodies from convalescent or vaccinated individuals, a small but convincing number of cross-neutralizing antibodies with similar epitope specificity have also been found ^39, 49–60^. The epitopes recognized by the G1, G2, and G3 nanobodies are therefore not only the viable targets in alpaca but also in humans. In particular, the G1 and G2 nanobodies would fall into the inner face antibodies exemplified by CR3022, H014, S2X259, COVA1-16, CV2-75, and C1c-A3 whereas the G3 nanobodies with the escarpment face antibody 47D11 (Figure S5)^61^. Additional cross-neutralizing SARS-CoV-2 and SARS-CoV-1 antibodies targeting to other faces of RBD have also been identified such as those to the cliff face (S2H97 and 6D60), top face (S2X146), and outer face (S309 and BG10-19) (Figure S5). Recently, a human antibody with broad reactivity to human beta-coronaviruses has been isolated that targets the conserved S2 stem-helix, raising the possibility of development of pan-β-CoV therapies and vaccines ^62^. More importantly, these cross-neutralizing and pan-β-CoV antibodies can be substantially boosted by mRNA vaccines, particularly in individuals with pre-existing SARS-CoV-1 or SARS-CoV-2 infection ^63–65^. Comprehensive characterization of these antibodies will provide us with deeper insights into their ontogeny and potential ways of inducing broader and more effective protection against the circulating and future variants.

Taken together, identification of 3-2A2-4 in this work and broadly neutralizing human antibodies elsewhere has revealed the existence of highly conserved and vulnerable sites on the RBD and spike trimer that could potentially be explored to trigger broader and more protective immune responses. This would require preferential exposure of these conserved regions while minimizing the receptor-binding motif that was predominantly recognized during natural infection and vaccination. While to achieve this goal would be expected a huge challenge, recent advances in structure-based vaccine design through understanding of antigen and antibody interaction will undoubtedly provide new conceptual and technological toolbox for this highly anticipated outcome. Identification of 3-2A2-4, as well as other broadly neutralizing antibodies, represent the first but a crucial step for us to achieve the ultimate goal of developing universal vaccine against all SARS-CoV-2 variants including Omicron BA.1 and BA. 2, and beyond.

Given the nature of immune response in alpaca is likely different from that in human, the antibody responses and the epitopes recognized by the nanobodies may not be exactly reflective of that in humans. Furthermore, our nanobody protection experiments were performed exclusively in K18-hACE2 mice, which are inherently different in responses to infection of different SARS-CoV-2 variants. Omicron resulted in relatively milder disease and lower viral replication in lungs than Delta. The protection efficacies against the two variants should therefore not be compared. Future studies in NHP and humans would be highly desirable to verify and validate the protection results.

## Acknowledgements

This study was funded by the National Key Plan for Scientific Research and Development of China (2020YFC0848800, 2020YFC0849900, 2021YFC0864500, 2020YFC0861200 and 2021YFC2300104), the National Natural Science Foundation (81530065, 81661128042, 9216920007 91442127, 32000661 and 32171202), Beijing Municipal Science and Technology Commission (D171100000517001, D171100000517003 and Z201100005420019), the Science and Technology Innovation Committee of Shenzhen Municipality (202002073000002 and JSGG20200807171401008), Beijing Advanced Innovation Center for Structural Biology, Tsinghua University Scientific Research Program (20201080053 and 2020Z99CFG004), Tencent Foundation, Shuidi Foundation, TH Capital, and the National Science Fund for Distinguished Young Scholars (82025022), Singapore National Medical Research Council Centre Grant Program (CGAug16M009), NUHSRO/2020/066/NUSMedCovid/01/BSL3 Covid Research Work, NUHSRO/2020/050/RO5+5/NUHS-COVID/4, Singapore Ministry of Health MOH-COVID19RF2-0001. We thank the SSRF BL02U1 and BL18U1 beam line for data collection and processing. We thank the X-ray crystallography platform of the Tsinghua University Technology Center for Protein Research for providing the facility support.

## Author contributions

L.Z., X.W. and J.C. conceived and designed the study. M.L, Y.R., Z.A. and B.C. performed most of the experiments with assistance from Y.L., Q.L. J.H., Y.Y., Y.W., J.C., S.S. J.G., X.S. and Q.Z. B.C. immunized the alpaca and constructed the yeast library. M.L., Y.L, Q.L, J.H., and Y.Y. isolated nanobodies and performed all evaluations. Y.R., J.C. and J.G. solved and analyzed crustal structure of nanobody and RBD complexes. Z.A., Y.W. and J.C. performed the nanobody protection experiment in K18-hACE2 mice. Z.Y. conducted the sequence analysis. L.C. conducted the live SARS-CoV-2 neutralization assay. R.W. constructed the pseudoviruses of SARS-CoV-2 and variants. M.L. and Y.R. had full access to data in the study, generated figures and tables, and take responsibility for the integrity and accuracy of the data presentation. L.Z., X.W, M.L. and Y.R. wrote the manuscript. All authors reviewed and approved the final version of the manuscript.

## METHOD DETAILS

### Cell lines

HEK293T cells (ATCC, CRL-3216) and HeLa cells expressing hACE2 were kindly provided by Dr. Qiang Ding at Tsinghua University. Both of these cell lines were maintained at 37°C in 5% CO_2_ in Dulbecco’s minimal essential medium (DMEM) containing 10% (v/v) heat-inactivated fetal bovine serum (FBS) and 100 U/mL penicillin–streptomycin. FreeStyle 293F cells (Thermo Fisher Scientific, R79007) were maintained at 37°C in 5% CO_2_ in SMM 293-TII expression medium (Sino Biological, M293TII). Sf9 cells (ATCC) were maintained at 27°C in Sf-900 II SFM medium. Hi5 cells (ATCC) were maintained at 27°C in SIM HF medium.

### Expression and purification of recombinant proteins

The genes encoding the ectodomain of S trimer and S2 trimer of prototype SARS-CoV-2 Wuhan-Hu-1 strain (GenBank: MN908947.3) were constructed as previously reported (Ruoke Wang, Immunity 2021). Both S trimer (residues M1-Q1208) and S2 trimer (residues S686-Q1208) contained proline substitutes at residue 986 and 987, a foldon trimerization motif, and a strep tag at C-terminal. Additional “GSAS” substitution at furin cleavage site (residues 682-685) was also introduced into the S trimer construct to improve overall stability. Both S trimer and S2 trimer were expressed in the FreeStyle 293F cells and purified by Strep-Tactin Sepharose (IBA Lifesciences) followed by gel filtration chromatography (GE Healthcare). The S trimer of SARS-CoV-1, RBD of SARS-CoV-1 and SARS-CoV-2 prototype, and the N-terminal peptidase domain of human ACE2 were expressed using the Bac-to-Bac Baculovirus Expression System (Invitrogen, Carlsbad, CA, USA) and purified by Strep-Tactin Sepharose (IBA Lifesciences) followed by gel filtration chromatography (GE Healthcare), as previously reported ^66^. Briefly, the RBD of SARS-CoV-2 wildtype (residues Arg319–Lys529) with an N-terminal gp67 signal peptide for secretion and a C-terminal 6×His tag for purification was expressed using Hi5 cells and purified by Ni-NTA resin followed by gel filtration chromatography (GE Healthcare) in HBS buffer (10 mM HEPES, pH 7.2, 150 mM NaCl). The RBD of SARS-CoV-1 (residues Arg306–Leu515) and N-terminal peptidase domain of human ACE2 (residues S19–D615) were expressed and purified by the same protocol as used for the RBD of SARS-CoV-2 wildtype (see above). The SARS-CoV-2 NTD protein was purchased from SinoBiological (40591-V49H).

### Immunization of alpaca, construction of yeast display VHH library, and isolation of VHH yeasts specific for SARS-CoV-2 and SARS-CoV-1 spikes

The animal experiment protocol involving immunization, collection of blood samples, and construction of VHH library was approved by IACUC at NBbiolab, Inc. in Chengdu, China. The immunization procedure involved three-time subcutaneous injections of 200 μg recombinant RBD of SARS-CoV-2 in Freund adjuvant, one-time subcutaneous injection of 10^11^ viral particle AdC68-19S vaccine expressing the wildtype S trimer ^67^, and two-time subcutaneous injections of 200 μg recombinant SARS-CoV-2 S-2P protein in Freund adjuvant. Seven days after the last immunization, blood samples were collected to isolate peripheral blood lymphocytes and plasma. Total RNA was extracted from the peripheral blood lymphocytes and used as the template for the first strand cDNA synthesis, using oligo dT as a primer. VHH sequences were amplified by PCR, cloned into a yeast surface display vector pYD1, and introduced into the electrocompetent EBY100 cells. The yeast library was first grown in SDCAA media at 30 °C for 48 h. At the exponential growth phase, the yeast library was transferred to SGCAA media for induction of VHH expression at 20 °C for 36 h. The yeast clones displaying VHHs specific to SARS-CoV-2 or SARS-CoV-1 spikes were enriched by one round of MACS biopanning followed by additional round of FACS biopanning. Specifically, induced yeast library was collected and incubated with 100 nM SARS-CoV-2 S-2P protein or SARS-CoV-1 S protein on ice for 30 min. The yeast library was washed with cold PBS+1%FBS for three times and incubated with streptavidin microbeads on ice for 10 min. The mixture was passed through LS column and the S protein-positive yeasts were harvested for further culture and induction of VHH expression. Once again, induced yeasts were incubated with 100 nM SARS-CoV-2 S-2P protein or SARS-CoV-1 S protein on ice for 30 min. After extensive wash with cold PBS+1%FBS, the yeast clones were incubated with HA-Tag (6E2) mouse monoclonal antibody conjugated with Alexa Fluor® 488 (1:100 dilution) and eBioscience™ streptavidin conjugated with PE Conjugate (1:200 dilution) on ice for 30 min. The yeast clones were washed three times with cold PBS+1%FBS before analyzed by FACS using Aria II (BD Biosciences). Positive yeast clones were sorted and used for nanobody cloning.

### Molecular cloning and expression of VHH

The sequences encoding various VHHs were amplified from the sorted yeast clones and inserted into two different expression vectors depending on the subsequent study objectives. One was to express VHH in conjunction with human IgG1 Fc fragment to evaluate binding and neutralizing activity of VHHs. The other was to express VHH with a 6xHis tag for crystal structural analysis. For the former, VHH genes were cloned into the multiple cloning sites of pMD18T containing the upstream CMV promoter, the secretory signal sequence from the mouse Ig heavy chain, and the downstream human IgG1 Fc gene fragment and SV40 poly (A) signal sequence. For the latter, selected VHH genes were cloned into pVRC8400 vector with a 6xHis tag. Expression and production of nanobodies were conducted by transfecting the expression vectors into the HEK293F cells using polyethyleneimine (PEI) (Polysciences). After approximately 96 h, nanobodies containing human IgG1 Fc in the culture supernatant were captured by AmMag Protein A Magnetic Beads (Genscript L00695) and eluted by Glycine pH 3.0. Nanobodies with 6xHis tag were captured by Ni-NTA Agaose (QIAGEN) and eluted by 500 nM Imidazole. All nanobodies were purified by gel-filtration chromatography with Superdex 200 High-Performance column (GE Healthcare). The exact concentration was determined by nanodrop 2000 Spectrophotometer (Thermo Scientific).

### Production of SARS-CoV-2 and sarbecovirus pseudoviruses

Pseudoviruses carrying the full-length spike envelope of SARS-CoV-2 and sarbecoviruses were generated as previously reported ^6^. Specifically, human immunodeficiency virus backbones expressing firefly luciferase (pNL4-3-R-E-luciferase) and pcDNA3.1 vector encoding either SARS-CoV-2 or sarbecovirus spike proteins were co-transfected into the HEK-293T cells (ATCC). Forty-eight hours later, pseudoviruses in the viral supernatant were collected, centrifuged to remove cell lysis, and stored at −80°C until use. The wildtype pseudovirus used throughout the analysis was the prototype strain (GenBank: MN908947.3) with a D614G mutation (WT D614G). The Alpha variant (Pango lineage B.1.1.7, GISAID: EPI_ISL_601443) included a total of 9 reported mutations in the spike protein (69-70del, 144del, N501Y, A570D, D614G, P681H, T716I, S982A and D1118H). The Beta variant (Pango lineage B.1.351, GISAID: EPI_ISL_700450) included 10 identified mutations in the spike such as L18F, D80A, D215G, 242-244del, S305T, K417N, E484K, N501Y, D614G and A701V. The Gamma variant (Pango lineage P.1, GISAID: EPI_ISL_792681) had 12 reported mutations in the spike including L18F, T20N, P26S, D138Y, R190S, K417T, E484K, N501Y, D614G, H655Y, T1027I and V1176F. The Delta variant (Pango lineage B.1.617.2, GISAID: EPI_ISL_1534938) included 10 reported mutations in the spike such as T19R, G142D, 156-157del, R158G, A222V, L452R, T478K, D614G, P681R, D950N. The Omicron BA.1 variant (Pango lineage BA.1, GISAID: EPI_ISL_6752027) was constructed with 34 mutations in the spike such as A67V, 69-70del, T95I, G142D, 143-145del, 211del, L212I, ins214EPE, G339D, S371L, S373P, S375F, K417N, N440K, G446S, S477N, T478K, E484A, Q493R, G496S, Q498R, N501Y, Y505H, T547K, D614G, H655Y, N679K, P681H, N764K, D796Y, N856K, Q954H, N969K, and L981F. The Omicron BA.2 variant (Pango lineage BA.2, GISAID: EPI_ISL_8515362) was constructed with 29 mutations in the spike such as T19I, 24-26del, A27S, G142D, V213G, G339D, S371F, S373P, S375F, T376A, D405N, R408S, K417N, N440K, S477N, T478K, E484A, Q493R, Q498R, N501Y, Y505H, D614G, H655Y, N679K, P681H, N764K, D796Y, N969K and Q954H. The Kappa variant (Pango lineage B.1.617.1, GISAID: EPI_ISL_1384866) was constructed with 8 mutations in the spike such as T95I, G142D, E154L, L452R, E484Q, D614G, P681R and N1071H. The Eta variant (Pango lineage B.1.525, GISAID: EPI_ISL_ 2885901) included 8 mutations in the spike such as Q52R, A67V, 69-70del, Y144del, E484K, D614G, Q677H and F888L. The Iota variant (Pango lineage B.1.526, GISAID: EPI_ISL_ 2922249) was constructed with 6 mutations in the spike such as L5F, T95I, D253G, E484K, D614G and A701V. The Lambda variant (Pango lineage C.37, GISAID: EPI_ISL_ 2930541) was constructed with 8 mutations in the spike such as G75V, T76I, R246N, 247-253del, L452Q, F490S, D614G and T859N. The Epsilon variant (Pango lineage B.1.429, GISAID: EPI_ISL_ 2922315) was constructed with 4 mutations in the spike such as S13I, W152C, L452R and D614G. The variant with Pango lineage A23.1, GISAID: EPI_ISL_ 2690464) was constructed with 4 mutations in the spike such as F157L, V367F, Q613H and P681R. The G339D, S371L, S371F, S373P, T376A, R408S and S371L+D373P+S375F were constructed with a D614G mutation based on the WT strain. For sarbecoviruses, the cDNAs encoding the SARS-CoV-1 S glycoprotein (NCBI Accession NP_828851.1), Pangolin CoV GD spike (GenBank: QLR06867.1), Pangolin CoV GX spike (GenBank: QIA48614.1), Bat SARS-like coronavirus WIV16 (GenBank: ALK02457.1), Bat SARS-like RaTG13 spike (GenBank: QHR63300.2), Bat SARS-like ZC45 (GenBank: AVP78031.1), Bat SARS-like ZXC21 (GenBank: AVP78042.1) and Bat SARS-like coronavirus RsSHC014 (GenBank: AGZ48806.1) were synthesized with codons optimized for protein expression (Genwiz Inc., China) and verified by sequencing.

### Neutralization activity of nanobodies against pseudoviruses and live SARS-CoV-2

Neutralization activity of nanobodies were determined using SARS-CoV-2 pseudovirus and authentic live virus as previously reported ^31^. Nanobodies were 3-fold serially diluted in 96-well cell culture plates, mixed with SARS-CoV-2 pseudovirus, and incubated at 37°C for 1 h. HeLa-ACE2 cells were then added to the mixture of nanobody-pseudovirus, incubated at 37°C for additional 48 h, and lysed for measuring luciferase-activity. The IC50 values were calculated based on the reduction of 50% relative light units (Bright-Glo Luciferase Assay Vector System, Promega, USA) compared to the virus-only control, using Prism 8.0 (GraphPad Software Inc., USA).

For the authentic live virus assay, we used the focus reduction neutralization test (FRNT) performed in a certified BSL3 facility at Shenzhen Third People’s Hospital, China. Briefly, serial dilutions of nanobodies were mixed with different authentic SARS-CoV-2 and incubated for 1 h at 37°C. The mixtures were then transferred to 96-well plates seeded with Vero E6 cells and incubated for 1 h at 37°C. After changing the medium, the plates were incubated at 37°C for an additional 24 h. The cells were then fixed, permeabilized, and incubated with cross-reactive rabbit anti-SARS-CoV-N IgG (Sino Biological, Inc., China) for 1 h at room temperature before adding an HRP-conjugated goat anti-rabbit IgG antibody (Jackson ImmunoResearch, USA). The reactions were developed using KPL TrueBlue peroxidase substrate (Seracare Life Sciences Inc., USA). The number of SARS-CoV-2 foci was quantified using an EliSpot reader (Cellular Technology Ltd. USA). The authentic SARS-CoV-2 used in the assay including the WT, Alpha, Beta, and Delta were isolated locally. Their whole genome sequences have been deposited into China National Center for Bioinformation, with accession numbers GWHXXXX00000000 for WT strain, GWHBFWX01000000 for Alpha strain, GWHBDSE01000000 for Beta strain, and GWHBFWZ01000000 for Delta strain, which are publicly accessible at https://ngdc.cncb.ac.cn/gwh.

### Phylogenetic tree and genetic analysis of nanobodies

Neighbor-joining phylogenetic trees were generated using MEGA version 10.1.8 with 1000 bootstrap replicates ^68^. The IMGT/V-QUEST program (http://www.imgt.org/IMGT_vquest/vquest) was used to analyze the germline gene and the loop lengths of complementarity determining region 3 (CDR3) of each nanobody. Chord diagrams showing the germline gene usages and V/J gene pairing were analyzed and presented by the R package circlize version 0.4.13 ^69^. The width of linking arc is proportional to the number of nanobodies identified. Sequence logo were plotted using Python package Logomaker ^70^.

### Nanobody binding kinetics and competition with ACE2 measured by SPR

The binding kinetics of nanobodies to SARS-CoV-2 RBD were analyzed using SPR (Biacore 8K; GE Healthcare). The nanobodies were captured by the ProA sensor chip and serial dilutions of SARS-CoV-2 RBD were then flowed through at a rate of 30 μL/min in PBS buffer (with 0.05% Tween20) for 120s with a dissociation time 3600s. The resulting data were fitted to a 1:1 binding model using the Biacore 8K Evaluation software (GE Healthcare). To determine the levels of nanobody competition with the human ACE2 and VHH72 of known epitope specificity, the prototype SARS-CoV-2 RBD was immobilized to a CM5 sensor chip via the amine group for a final RU around 250. In the first round, nanobodies (1 μM) or VHH72 (1 μM) were injected onto the chip for 120 s to reach the steady state for binding. In the second round, nanobodies (1 μM) or VHH72 (1 μM) were injected onto the chip for 120 s followed by injection of ACE2 (2 μM) or nanobodies (1 μM) for additional 120 s. In the third round, running buffer was injected for 120 s followed by injection of ACE2 (2 μM) or nanobodies (1 μM) for another 120 s. The sensorgrams of the three rounds were aligned from 120 to 240 s in Biacore 8K Evaluation software (GE Healthcare). The blocking efficacy was determined by a comparison of the response units with and without prior antibody injection.

### Crystallization and data collection

To obtain the complex of RBD bound to nanobodies, RBD was incubated with each nanobody for 1 h on ice in HBS buffer. The mixture was then purified using gel filtration chromatography. Fractions containing the complex were pooled and concentrated to 10 mg ml−1. Crystals were successfully grown at room temperature in sitting drops, over wells containing 0.2 M Ammonium sulfate, 0.1 M Bis-Tris pH 4.4, 21% w/v Polyethylene glycol 3,350 for Nb70-SARS-CoV-1 RBD, 0.15M DL-Malic acid pH 7.0, 20% w/v Polyethylene glycol 3,350 for Nb70-1F11 fab-SARS-CoV-2 RBD, 0.2 M Ammonium sulfate, 0.1M Bis Tris pH 5.5, 25% w/v Polyethylene glycol 3,350 for 1-2C7-SA RBD, 0.2 M Ammonium formate, pH 6.6, 20% w/v Polyethylene glycol 3,350 for 3-2A2-4-SARS-CoV-2 RBD. Crystals were collected, soaked briefly in 0.2 M Ammonium sulfate, 0.1 M Bis-Tris pH 4.4, 21% w/v Polyethylene glycol 3,350 and 20% glycerol for Nb70-SARS-CoV-1 RBD, 0.15 M DL-Malic acid pH 7.0, 20% w/v Polyethylene glycol 3,350 and 20% glycerol for Nb70-1F11 fab-SARS-CoV-2 RBD, 0.2 M Ammonium sulfate, 0.1 M Bis-Tris pH 5.5, 25% w/v Polyethylene glycol 3,350 and 20% glycerol for 1-2C7-SA RBD, 0.2 M Ammonium formate, pH 6.6, 20% w/v Polyethylene glycol 3,350 and 20% glycerol for 3-2A2-4-SARS-CoV-2 RBD and were subsequently flash-frozen in liquid nitrogen. Diffraction data were collected at 100 K and a wavelength of 0.987 Å at the BL18U1 beam line for Nb70-SARS-CoV-1 RBD, Nb70-1F11 fab-SARS-CoV-2 RBD and 1-2C7-SA RBD, 100 K and a wavelength of 1.07180 Å at the BL02U1 beam line for 3-2A2-4-SARS-CoV-2 RBD of the Shanghai Synchrotron Research Facility. Diffraction data were processed using the HKL3000 software (PMID:16855301) and the data-processing statistics are listed in Table S1.

### Data availability

The coordinates and structure factors for the 1-2C7-SARS-CoV-2 SA-RBD, Nb70-SARS-CoV-2 WT-RBD-P2C-1F11, Nb70-SARS-CoV-1 WT-RBD, 3-2A2-4-SARS-CoV-2 WT-RBD complex were deposited in Protein Data Bank with accession code 7X2M, 7X2K, 7X2J, 7X2L.

### Structure determination and refinement

The structure was determined using the molecular replacement method with PHASER in the CCP4 suite (PMID: 19461840). Density map improvement by updating and refinement of the atoms was performed with ARP/wARP26 (PMID: 18094467). Subsequent model building and refinement were performed using COOT (PMID: 15572765) and PHENIX (PMID: 12393927), respectively. Final Ramachandran statistics: 96.54% favored, 3.46% allowed and 0.00% outliers for the final Nb70-SARS-CoV-1 RBD structure, 97.28% favored, 2.59% allowed and 0.14% outliers for the final Nb70-1F11 fab-SARS-CoV-2 RBD structure, 97.15% favored, 2.85% allowed and 0.00% outliers for the final 1-2C7-SA RBD structure and 93.89%favored, 5.79% allowed and 0.32% outliers for the final 3-2A2-4-SARS-CoV-2 RBD structure. The structure refinement statistics are listed in Table S1. All structure figures were generated with ChimeraX and Pymol (PMID: 28158668).

### Nanobody protection in K18-hACE2 mice

Animal experiments were conducted in a Biosafety Level 3 (BSL-3) facility in accordance with the National University of Singapore (NUS) Institutional Animal Care and Use Committee (IACUC) (protocol no. R20-0504), and the NUS Institutional Biosafety Committee (IBC) and NUS Medicine BSL-3 Biosafety Committee (BBC) approved SOPs. Eight-week-old female K18-hACE2 transgenic mice (InVivos Ptd Ltd, Lim Chu Kang, Singapore) were used for this study. The mice were housed and acclimatized in an ABSL-3 facility for 72 h prior to the start of the experiment. K18-hACE2 transgenic mice were subjected to pretreatment of nanobody 3-2A2-4 (10 mg/kg) delivered through intraperitoneal injection a day prior to infection. The viral challenge was conducted through intranasal delivery in 25 μl of either 1.7×10^3^ PFU of the infectious SARS-CoV-2 Omicron or 10^3^ PFU of Delta variant. Baseline body weights were measured prior to infection and monitored daily by two personnel post-infection for the duration of the experiment. Mice were euthanized when their body weight fell below 75% of their baseline body weight. To assess the viral load, mice from each experimental group were sacrificed 3 days post inoculation, with lung tissues harvested. Each organ was halved for the plaque assay and histology analysis, respectively. Tissues were homogenized with 0.5 mL DMEM supplemented with antibiotic and antimycotic (Gibco, Waltham, MA, USA) and titrated in Vero E6 cells using plaque assays. For virus titer determination, supernatants from homogenized tissues were diluted 10-fold serially in DMEM supplemented with antibiotic and antimycotic. Of each serial diluted supernatant, 250 μl was added to Vero E6 cells into 12-well plates. After 1 h of incubation for virus adsorption, the inoculum was removed and washed once with PBS. About 1.2% microcrystalline cellulose (MCC)-DMEM supplemented with antibiotic and antimycotic overlay media was added to each well and incubated at 37°C, 5% CO_2_ for 72 h for plaque formation. The cells were then fixed in 10% formalin overnight and counterstained with crystal violet. The number of plaques was determined and the virus titers of individual samples were expressed in logarithm of PFU per organ. For histopathological analyses, lung lobes were fixed in 3.7% formaldehyde solution prior to removal from BSL-3 containment. The tissues were routinely processed, embedded in paraffin blocks (Leica Surgipath Paraplast), sectioned at 4-μm thickness, and stained with H&E (Thermo Scientific) following standard histological procedures. For immunohistochemistry, sections were deparaffinized and rehydrated, followed by heat-mediated antigen retrieval, quenching of endogenous peroxidases and protein blocking. Sections were then covered with rabbit anti-SARS-CoV-2 N protein monoclonal antibody (Abcam; 1:1000) for 1 h at room temperature. Subsequently, sections were incubated with rabbit-specific HRP polymer (secondary antibody), visualized using chromogenic substrate DAB solution (Abcam), and counterstained with hematoxylin.

### Statical analysis

The technical and independent experiment replicates were indicated in the figure legends. Half-maximal inhibitory concentration (IC50) of nanobodies was calculated by the equation of four-parameter dose inhibition response using Graphpad Prism 8.0. The fold change of the variants relative to D614G in neutralization were calculated by simple division of respective IC50 values. In animal experiments, a two-tailed unpaired Mann-Whitney test was used to assess statistical significance. Statistical calculations were performed in GraphPad Prism 8.0. Differences with p-values less than 0.05 were considered to be statistically significant (**p < 0.01).

## SUPPLEMENTAL INFORMATION

**Figure S1, related to Figure 1.**
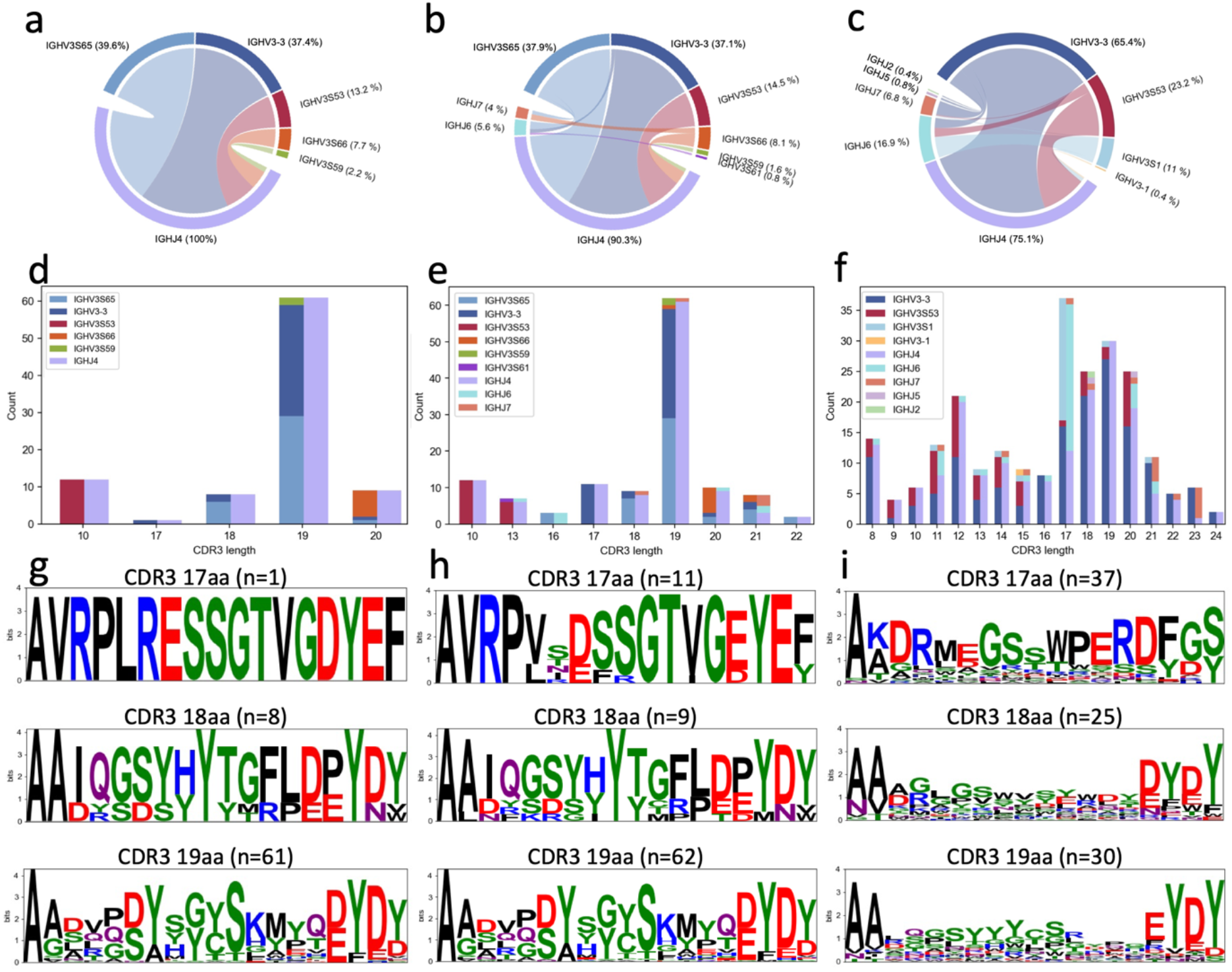
Genetic characterization and comparison between isolated and published nanobodies against SARS-CoV-2. **(a, b, c)** Chord diagrams comparing the V and J gene segments usage and pairing among the 91 cross-neutralizing, total 124 isolated, and published 237 nanobodies in the CoV-AbDab database. Each V and J segments are colored and indicated around the peripheral circle together with their percentage among the total number of nanobodies analyzed. V/J pairs are linked by colored arcs, and the size of which is proportional to the total number of nanobodies analyzed. **(d, e, f)** The bar plot showing the distribution and proportion of various CDR3 length among the 91 cross-neutralizing, total 124 isolated and 237 published nanobodies. The specific V and J gene usage associated with each CDR3 length are colored and shown. **(g, h, i)** Comparison of CDR3 logo sequence among the 91 cross-neutralizing, total 124 isolated, and 237 published nanobodies, analyzed separately for 17-residue (top), 18-residue (middle), and 19-reside (bottom) CDR3. The number of sequences analyzed for each CDR3 group are indicated.

**Figure S2, related to Figure 2.**
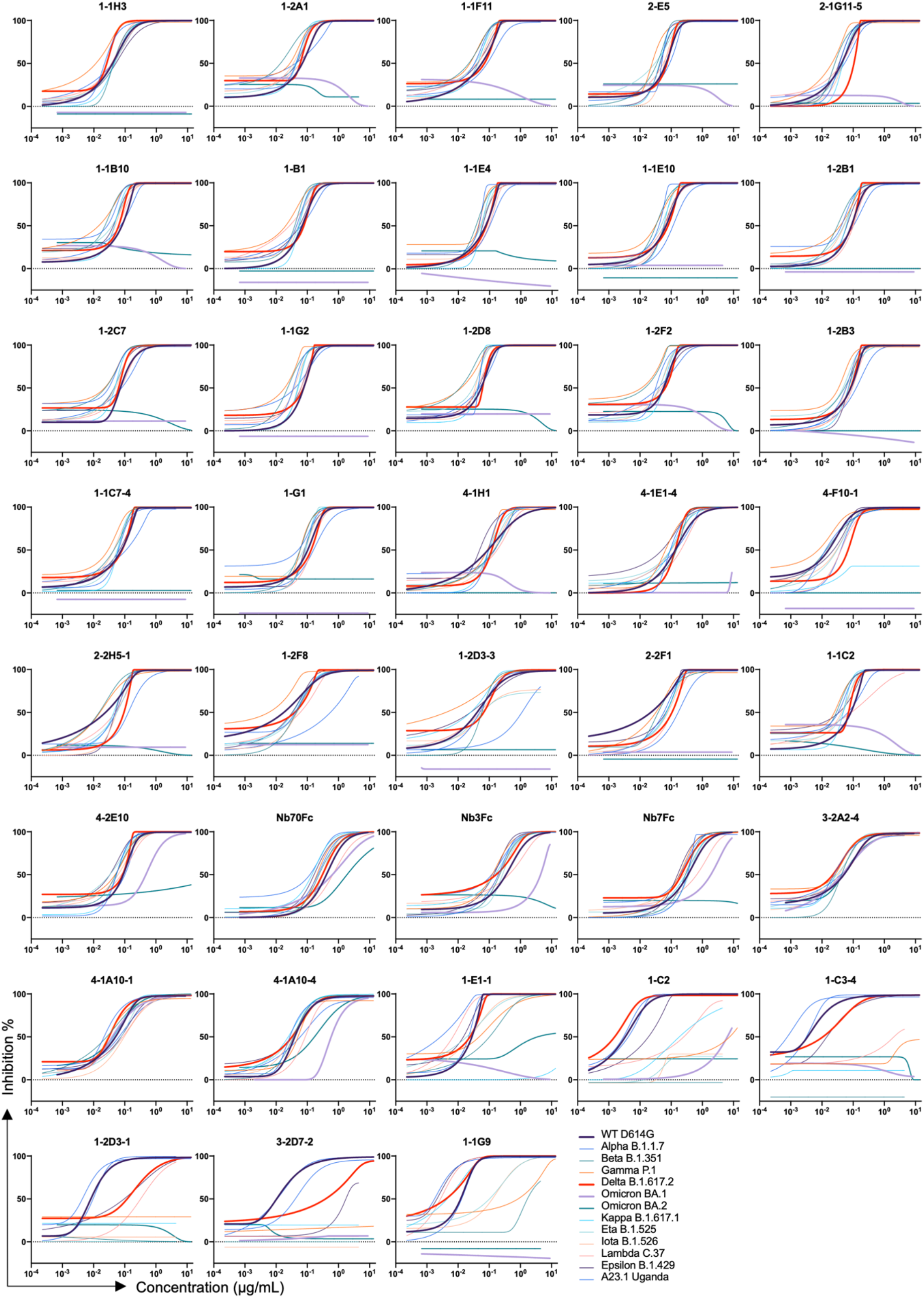
Neutralizing activity of isolated nanobodies against SARS-CoV-2 variants. Serial dilutions of each nanobody were evaluated against pseudoviruses carrying spike protein of prototype and variants of SARS-CoV-2. Neutralizing activity was defined as the percent reduction in luciferase activities compared to no antibody controls. Results presented are representatives of three independent experiments.

**Figure S3, related to Figure 2.**
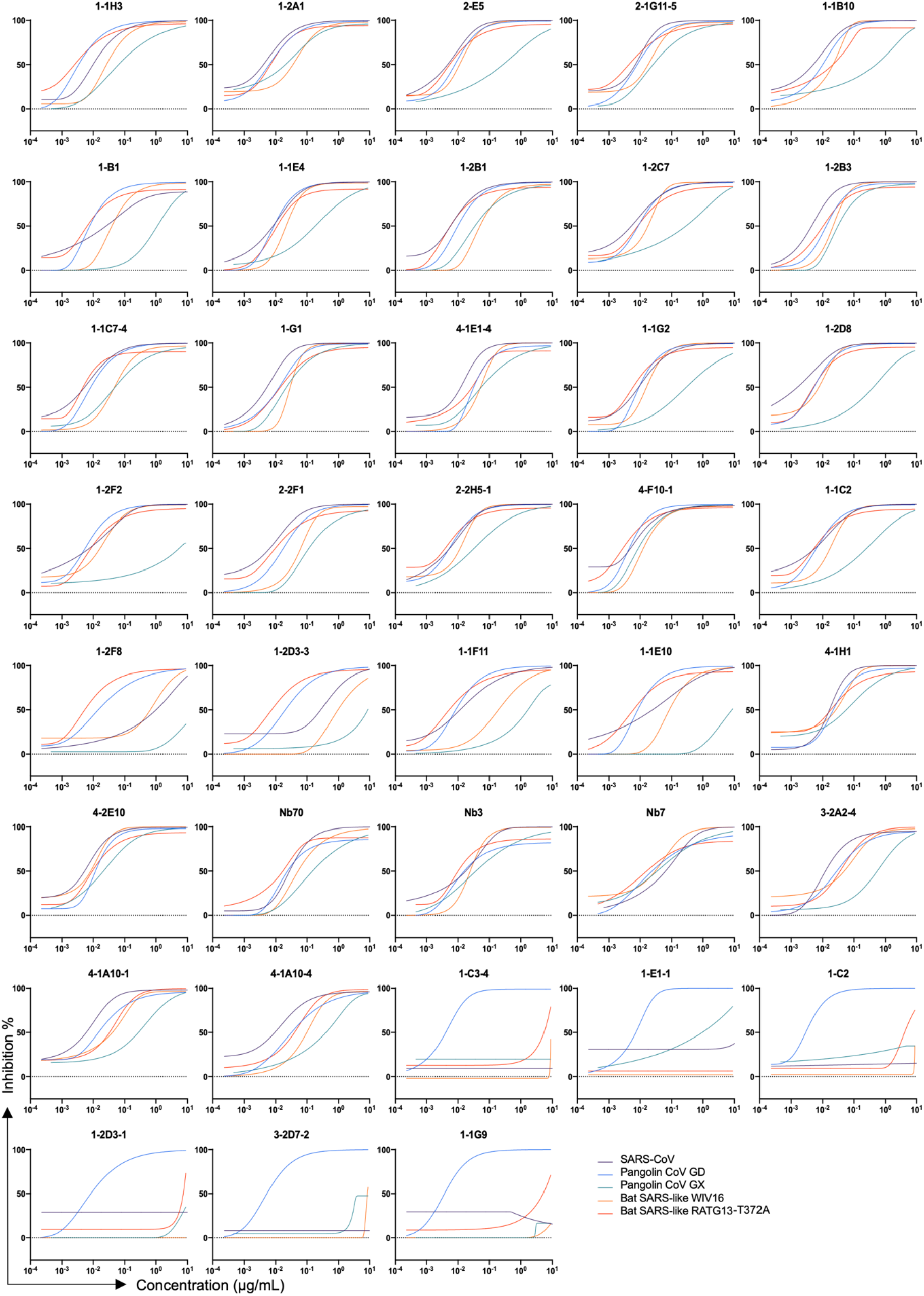
Neutralizing activity of isolated nanobodies against hACE2-dependent sarbecoviruses. Serial dilutions of each nanobody were tested against pseudoviruses carrying spike protein of various sarbecoviruses. Neutralizing activity was defined as the percent reduction in luciferase activities compared to no antibody controls. Results presented are representatives of three independent experiments.

**Figure S4, related to Figure 3.**
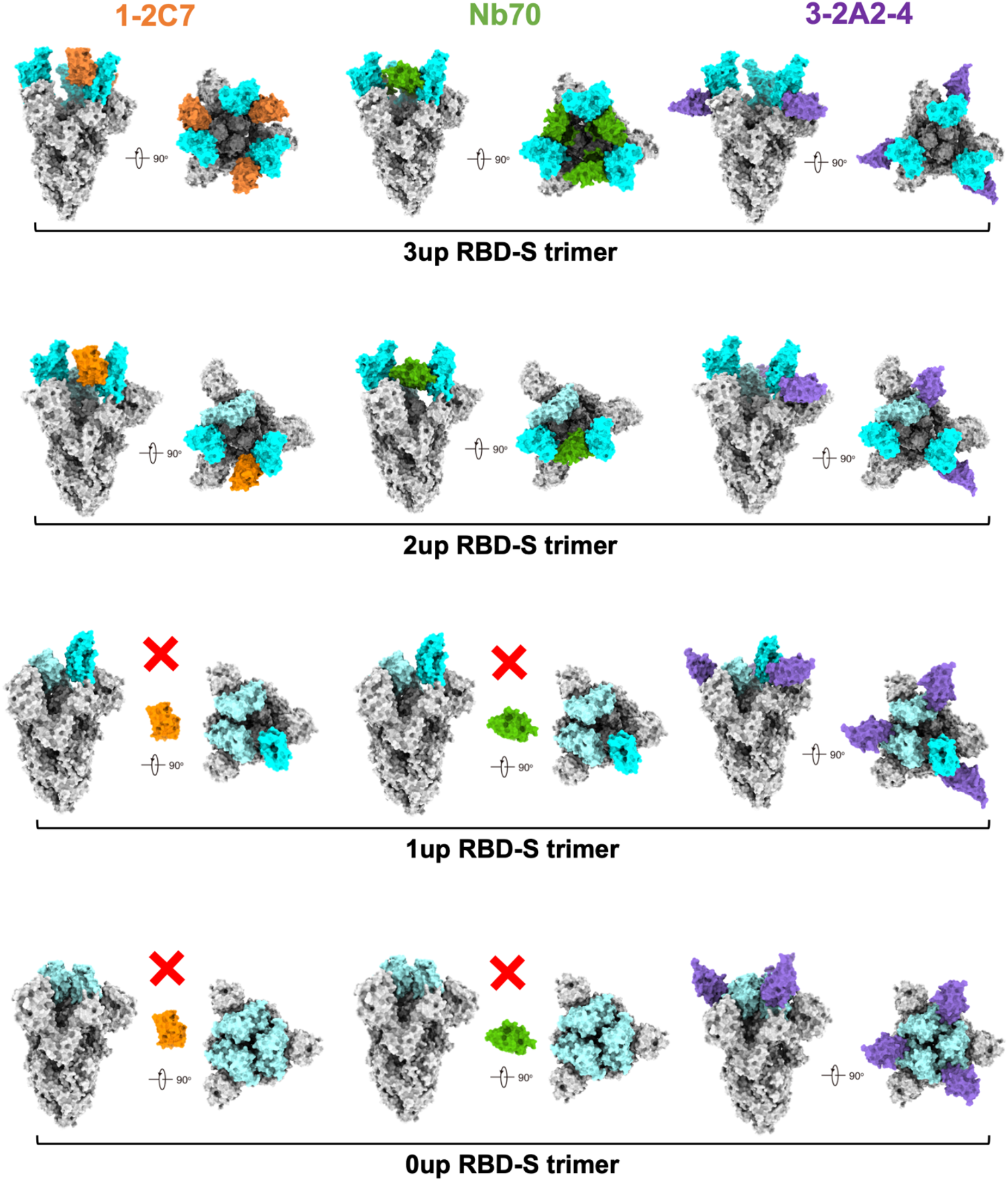
Proposed binding models of the three representative nanobodies to various RBD conformations in the context of prototype SARS-CoV-2 spike trimer. Crystal structures of 1-2C7, Nb70, and 3-2A2-4 bound to RBD are aligned to SARS-CoV-2 spike trimer in four different conformations: 1) 3up RBD-S trimer (PDB: 7KMS); 2) 2up RBD-S trimer (PDB: 7A93); 3) 1up RBD-S trimer (PDB: 6VYB); and 4) 0 up RBD-S trimer (PDB: 6VXX). The spike trimers are shown as a molecular surface, with up RBD colored in cyan, down RBD in light blue, NTD and S2 in grey. 1-2C7, Nb70, and 3-2A2-4 are colored in orange, green, and purple, respectively.

**Figure S5.**
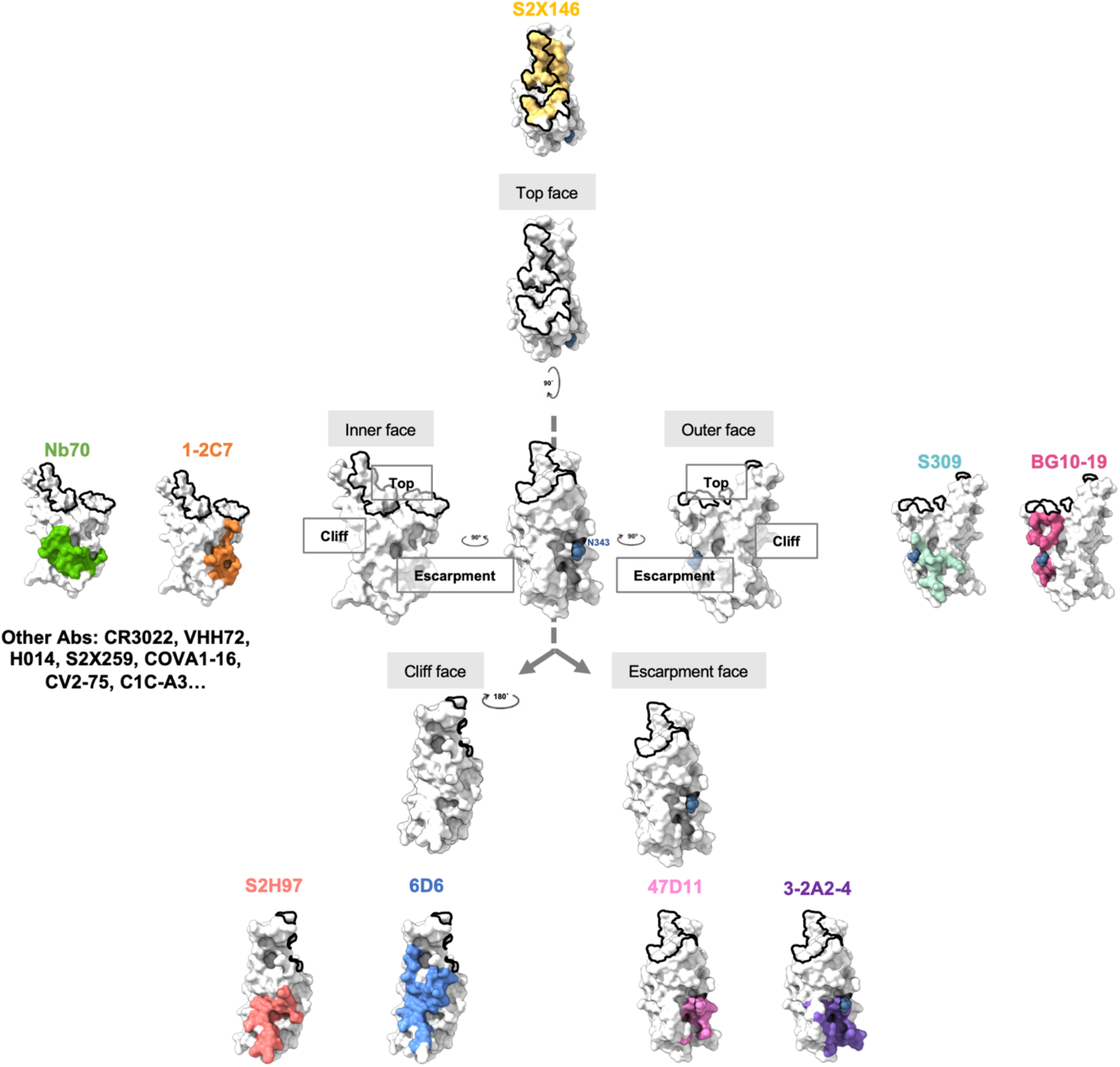
Structural illustration of SARS-CoV-1 and SARS-CoV-2 cross-neutralizing epitopes recognized by the three representative nanobodies and various published antibodies. Cross-neutralizing epitopes on the top face of RBD is recognized by S2X146, inner face by Nb70 and 1-2C7, outer face by S309 and BG10-19, cliff face by S2H97 and 6D6, and escarpment face by 47D11 and 3-2A2-4. The glycosylation site at position 343 (N343), conserved across the sarbecovirus subgenus, is colored in dark blue. The ACE2-binding site is outlined in black.

**Table S1, related to Figure 3.** Data collection and refinement statistics.

|  | Nb70-SARS-CoV-1<br>RBD | Nb70-1F11 Fab- SARS-CoV-<br>2 WT RBD | 1-2C7-SARS-CoV-2 SA<br>RBD | 3-2A2-4-SARS-CoV-2 WT<br>RBD |
| --- | --- | --- | --- | --- |
| **Data collection** |  |  |  |  |
| Space group | P3<sub>1</sub>21 | P2<sub>1</sub>2<sub>1</sub>2<sub>1</sub> | I23 | P4<sub>1</sub>2<sub>1</sub>2 |
| Cell dimensions |  |  |  |  |
| a, b, c (Å) | 108.413 108.413 94.119 | 94.629 95.47 104.139 | 143.619 143.619 143.619 | 89.168 89.168 129.154 |
| $\alpha$, $\beta$, $\gamma$ (°) | 90, 90, 120 | 90, 90, 90 | 90, 90, 90 | 90, 90, 90 |
| Resolution (Å) | 50-2.4(2.486-2.4) | 50-2.4(2.486-2.4) | 50-1.8(1.864-1.8) | 50-2.402 (2.488-2.402) |
| *R*<sub>merge</sub> | 0.135 (2.269) | 0.213 (1.249) | 0.172 (2.643) | 0.152 (2.555) |
| *I*/*sI* | 25.45 (1.10) | 17.18 (1.74) | 33.27 (1.57) | 23.43 (1.54) |
| Completeness (%) | 99.77 (99.84) | 99.54 (99.89) | 99.93 (100) | 97.37 (83.09) |
| Redundancy | 17.9 (14.3) | 12.2 (8.5) | 38.8(31.2) | 22.1 (20.9) |
| **Refinement** |  |  |  |  |
| Resolution (Å) | 33.24-2.4 | 45.62-2.4 | 25.39-1.8 | 28.33-2.402 |
| No. reflections | 25337 (2493) | 37381 (3676) | 45590 (4537) | 20419 (1695) |
| *R*<sub>work</sub>/*R*<sub>free</sub> | 19.98/23.78 | 19.53/24.36 | 17.99/20.97 | 22.96/28.74 |
| No. atoms |  |  |  |  |
| Protein | 375 | 745 | 323 | 315 |
| Ligand/ion | 28 | 14 | 39 | 49 |
| Water | 51 | 216 | 225 | 30 |
| B-factors |  |  |  |  |
| Protein | 64.96 | 43.35 | 44.17 | 69.22 |
| Ligand/ion | 103.73 | 52.62 | 67.20 | 69.84 |
| Water | 61.38 | 62.66 | 46.31 | 77.73 |
| R.m.s. deviations |  |  |  |  |
| Bond lengths(Å) | 0.008 | 0.008 | 0.007 | 0.010 |
| Bond angles (°) | 1.01 | 0.99 | 0.87 | 1.25 |
| Ramachandran |  |  |  |  |
| Favored (%) | 96.54 | 97.28 | 97.15 | 93.89 |
| Allowed (%) | 3.46 | 2.59 | 2.85 | 5.79 |
| Outliers (%) | 0.00 | 0.14 | 0.00 | 0.32 |

**Table S2, related to Figure 3.**
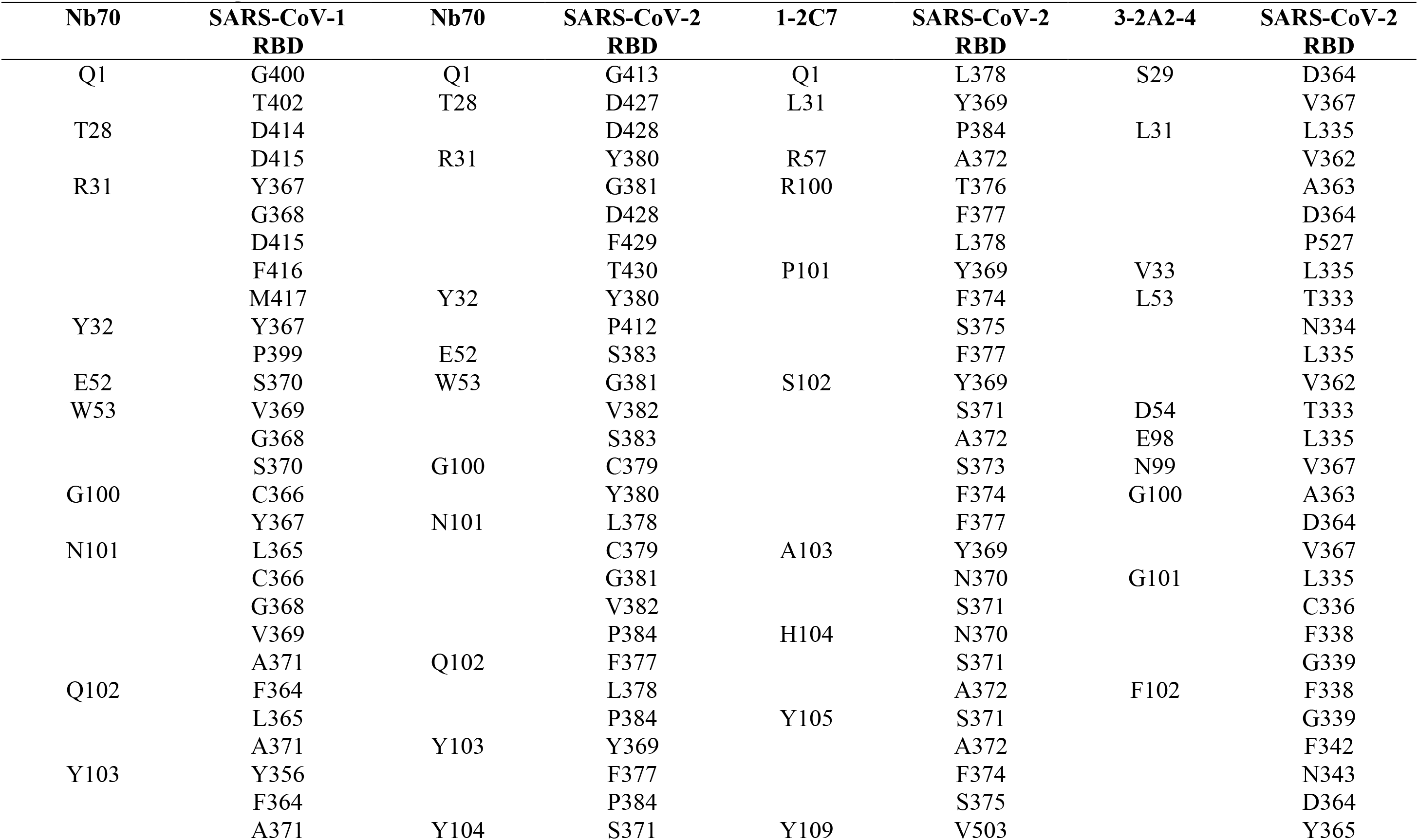
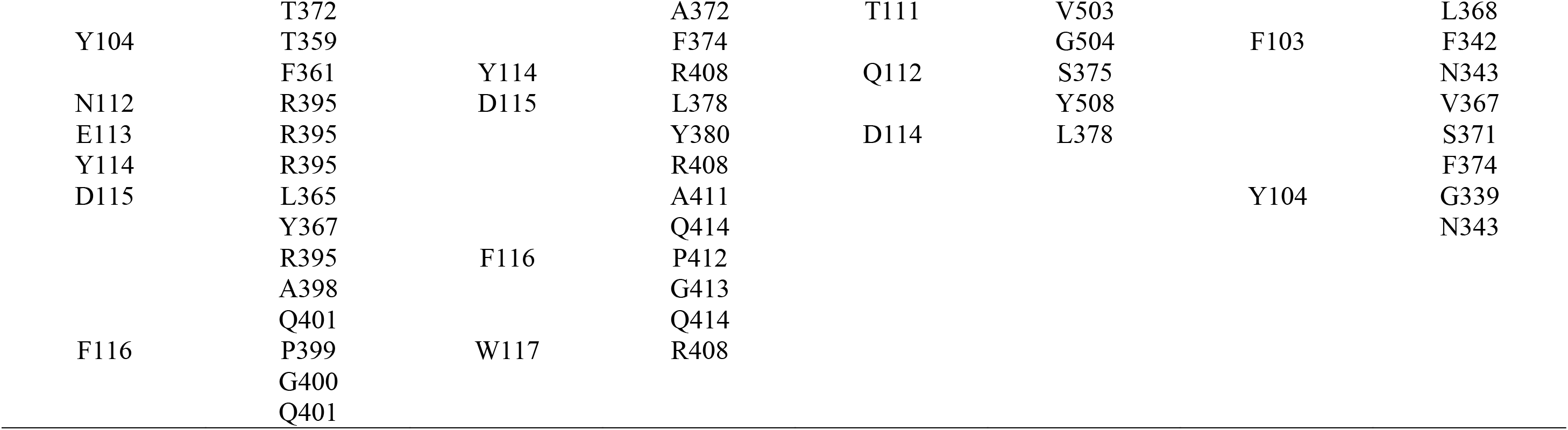
Contact residues of the Nbs-RBD interfaces.

